# Studying models of balancing selection using phase-type theory

**DOI:** 10.1101/2020.07.06.189837

**Authors:** Kai Zeng, Brian Charlesworth, Asger Hobolth

**Author notes:** Corresponding author: Department of Animal and Plant Sciences, University of Sheffield, Sheffield S10 2TN, United Kingdom.

## Abstract

Balancing selection (BLS) is the evolutionary force that maintains high levels of genetic variability in many important genes. To further our understanding of its evolutionary significance, we analyse models with BLS acting on a biallelic locus: an equilibrium model with long-term BLS, a model with long-term BLS and recent changes in population size, and a model of recent BLS. Using phase-type theory, a mathematical tool for analysing continuous time Markov chains with an absorbing state, we examine how BLS affects polymorphism patterns in linked neutral regions, as summarised by nucleotide diversity, the expected number of segregating sites, the site frequency spectrum, and the level of linkage disequilibrium (LD). Long-term BLS affects polymorphism patterns in a relatively small genomic neighbourhood, and such selection targets are easier to detect when the equilibrium frequencies of the selected variants are close to 50%, or when there has been a population size reduction. For a new mutation subject to BLS, its initial increase in frequency in the population causes linked neutral regions to have reduced diversity, an excess of both high and low frequency derived variants, and elevated LD with the selected locus. These patterns are similar to those produced by selective sweeps, but the effects of recent BLS are weaker. Nonetheless, compared to selective sweeps, non-equilibrium polymorphism and LD patterns persist for a much longer period under recent BLS, which may increase the chance of detecting such selection targets. An R package for analysing these models, among others (e.g., isolation with migration), is available.

---

**B**alancing selection refers to a type of natural selection that maintains genetic variability in populations (Fisher 1922; Charlesworth 2006; Fijarczyk and Babik 2015). Genes known to be under balancing selection are often involved in important biological functions. Examples include the major histocompatibility complex (MHC) genes in vertebrates (Spurgin and Richardson 2010), plant self-incompatibility genes (Castric and Vekemans 2004), mating-type genes in fungi (van Diepen *et al*. 2013), genes underlying host-pathogen interactions (Bakker *et al*. 2006; Hedrick 2011), inversion polymorphisms (Dobzhansky 1970), and genes underlying phenotypic polymorphisms in many different organisms (e.g., Johnston et al. 2013; Küpper et al. 2016; Kim et al. 2019). More recently, it has been proposed that a related process, known as associative overdominance, may play a significant role in shaping diversity patterns in genomic regions with very low recombination rates (Becher *et al*. 2020; Gilbert *et al*. 2020). These facts highlight the importance of studying balancing selection.

Understanding how balancing selection affects patterns of genetic variability is a prerequisite for detecting genes under this type of selection. The best studied models involve long-term selection acting at a single locus (Strobeck 1983; Hudson and Kaplan 1988; Takahata 1990; Takahata and Nei 1990; Vekemans and Slatkin 1994; Nordborg 1997; Takahata and Satta 1998; Innan and Nordborg 2003). It is well known that, in addition to maintaining diversity at the selected locus, long-term balancing selection increases diversity at closely linked neutral sites. This reflects an increased coalescence time for the gene tree connecting the alleles in a sample from the current population. When this tree is sufficiently deep, it is possible for the ages of the alleles to exceed the species’ age, leading to trans-species polymorphism. Furthermore, long-term balancing selection alters the site frequency spectrum (SFS) at linked neutral sites, causing an excess of intermediate frequency derived variants. These properties underlie most of the methods used for scanning large-scale genomic data for targets of balancing selection (Andres *et al*. 2009; Leffler *et al*. 2013; DeGiorgio *et al*. 2014; Bitarello *et al*. 2018; Cheng and DeGiorgio 2019; Siewert and Voight 2020).

A significant limitation of most previous studies is the assumption that the population is at statistical equilibrium under selection, mutation and genetic drift. In reality, most populations have experienced recent changes in population size. Our ability to analyse data from these populations is limited by the lack of an effective way of making predictions about the joint effects of demographic changes and balancing selection on patterns of genetic variability. Moreover, many cases of balancing selection involve variants that have only recently spread to intermediate frequencies, rather than having been maintained for periods much longer than the neutral coalescence time (e.g. Eanes 1999; Kwiatkowski 2005; Corbett-Detig and Hartl 2012). Indeed, several theoretical studies have suggested that adaptation may occur through the frequent emergence of short-lived balanced polymorphisms (Sellis *et al*. 2011; Connallon and Clark 2014). Because of their young age, there may not be sufficient time for the diversity patterns predicted for long-term balancing selection to emerge. As a result, targets of recent balancing selection are unlikely to be detected by existing methods. This may explain why genome scans have only reported a relatively small number of potential selection targets (Andres *et al*. 2009; Leffler *et al*. 2013; DeGiorgio *et al*. 2014; Bitarello *et al*. 2018; Cheng and DeGiorgio 2019).

Multiple authors have suggested that the emergence of a recent balanced polymorphism will generate diversity patterns that resemble those generated by incomplete selective sweeps caused by positive selection favouring a beneficial mutation (Charlesworth 2006; Sellis *et al*. 2011; Fijarczyk and Babik 2015). In fact, methods designed for detecting sweeps can identify these signals (e.g., Zeng *et al*. 2006). However, there is currently no theoretical framework for studying recent balanced polymorphism, which precludes a detailed comparison with incomplete selective sweeps. Acquiring this knowledge will help us devise methods for distinguishing between balancing selection and positive selection, which will in turn allow us to test hypotheses about the importance of balancing selection in adaptation.

We tackle these problems by using phase-type theory. Briefly, a phase-type distributed random variable describes the time until a continuous time Markov process enters one of its absorbing states. As an example, imagine that we have taken a sample of *n* alleles from the population. Going backwards, the time it takes for the process to reach the most recent common ancestor follows a phase-type distribution. Phase-type theory refers to a set of mathematical tools for analysing the properties (e.g., mean and variance) of this type of random variable (Bladt and Nielsen 2017). In a recent study, Hobolth *et al*. (2019) used a time-homogeneous version of the theory to study several population genetic models at statistical equilibrium. Here, we extend this approach by deriving several useful results under a time-inhomogeneous framework. We use the new theory to analyse three models of balancing selection: an equilibrium model of long-term balancing selection, a model with long-term balancing selection and changes in population size, and a model of recent balancing selection. The analysis of the last model is accompanied by a comparison with a comparable selective sweep model.

For each of these models, we calculate summary statistics that are useful for understanding the effects of selection on diversity patterns in nearby genomic regions. Specifically, for a sample of alleles collected from a linked neutral site, we obtain (1) the expected pairwise coalescence time (proportional to nucleotide diversity *π*), (2) the expected level of linkage disequilibrium (LD) between the selected locus and the focal neutral site, (3) the total branch length of the gene tree (proportional to the total number of segregating sites 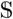), and (4) the site frequency spectrum (SFS). Our results extend previous studies of the equilibrium model by providing a unifying framework for obtaining these statistics. The analysis of the non-equilibrium models provides useful insights that can be used for devising new genome scan methods or parameter estimation methods. We conclude the study by discussing the usefulness of phase-type theory in population genetics.

## An equilibrium model of balancing selection

Consider a diploid, randomly mating population. The effective population size *N_e_* is assumed to be constant over time. An autosomal locus with two alleles *A*_1_ and *A*_2_ is under balancing selection. The intensity of selection is assumed to be sufficiently strong and constant over time that the frequencies of the two alleles remain at their equilibrium values indefinitely. Denote the equilibrium frequencies of *A*_1_ and *A*_2_ by 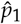 and 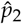, respectively 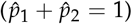. This set-up can accommodate any model of long-term balancing selection (with or without reversible mutation between *A*_1_ and *A*_2_), as long as it produces stable allele frequencies. A random sample of *n* alleles have been taken from a linked neutral locus. The recombination frequency between this locus and the selected locus is denoted by *r*. In the following four subsections, we use time-homogeneous phase-type theory to calculate the four statistics mentioned at the end of the Introduction. This introduces the methodology and notation, and sets the stage for extending the analysis to non-equilibrium models in later sections. A similar model has been investigated previously using different approaches (Strobeck 1983; Hudson and Kaplan 1988; Nordborg 1997). However, these do not provide analytical expressions for the SFS.

### The mean coalescence time for a sample size of two

An allele at the neutral locus is associated with either *A*_1_ or *A*_2_ at the selected site (i.e., a neutral allele is on the same haplotype as either *A*_1_ or *A*_2_). The sample is therefore in one of three possible states (Figure 1). In state 1, both alleles are associated with *A*_2_. In state 2, one allele is associated with *A*_1_, and the other is associated with *A*_2_. In state 3, both alleles are associated with *A*_1_. Take state 1 as an example. An allele currently associated with *A*_2_ was associated with *A*_1_ in the previous generation either because there was an *A*_1_ to *A*_2_ mutation during gamete production, or because the parent was an *A*_1_*A*_2_ heterozygote and there was a recombination event. Define *v*_21_ as the *backward* mutation rate (see Supplementary Text S.1). The first event occurs with probability *v*_21_, and the second event occurs with probability 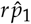. The probability that the focal allele becomes associated with *A*_1_ in the previous generation is 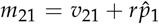. The two alleles in state 1 may share a common ancestor in the previous generation. Because the frequency of *A*_2_ is 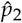, a total of 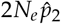 alleles were associated with *A*_2_ in the previous generation. The chance that the two alleles coalesce is 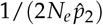.

**Figure 1.**
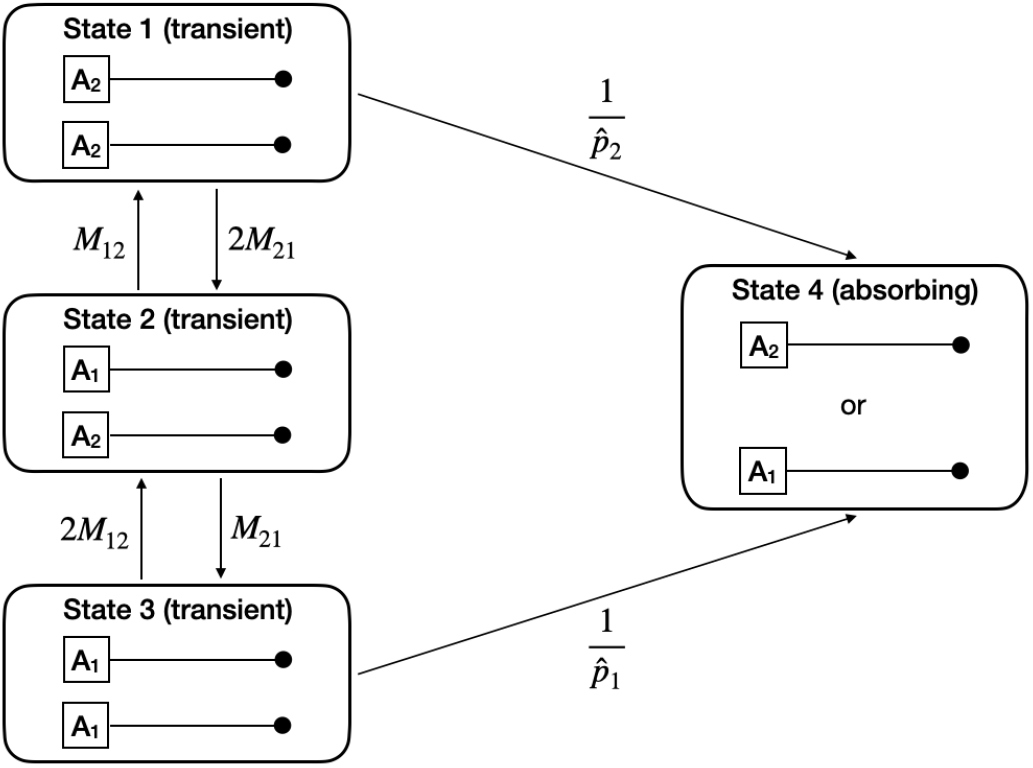
Transition rates between the states of the equilibrium balancing selection model for a sample size of two. A_1_ and A_2_ are the variants at the locus under balancing selection, with equilibrium frequencies 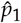 and 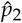, respectively. The *backward* mutation rate between *A_i_* and *A_j_* is *v_ij_* per generation. The thin horizontal lines represent haplotypes, and the neutral locus is represented by a black dot. The recombination frequency between the two loci is *r*. Time is scaled in units of 2*N_e_* generations. The rate at which a neutral allele associated with *A_i_* becomes associated with *A_j_* is 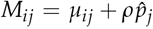, where *μ_ij_* = 2*N_e_v_ij_* and *ρ* = 2*N_e_r*. Two neutral alleles associated with *A_i_* coalesce at rate 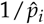.

Under the standard assumption that the probability of occurrence of more than one event in one generation is negligible, the probability that the two alleles in state 1 remain unchanged for *z* generations is:

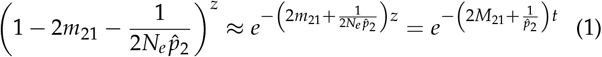

where 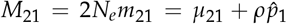, *μ*_21_ = 2*N_e_v*_21_, *ρ* = 2*N_e_r*, and *t = z*/(2*N_e_*).

We have scaled time in units of 2*N_e_* generations, and will use this convention throughout unless stated otherwise. Using this timescale, when in state 1, the waiting time to the next event follows an exponential distribution with rate parameter 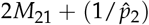. Given that an event has occurred, the probability that it is caused by one of the two alleles becoming associated with A_1_ is 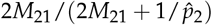, and the probability that it is caused by the coalescence of the two alleles is 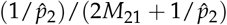. As illustrated in Figure 1, the first possibility moves the process from state 1 to state 2, whereas the second possibility terminates the process by moving it into the absorbing state where the most recent common ancestor (MRCA) is reached (state 4).

We can derive the transition rates between all four states of the process using similar arguments (Figure 1). This model is analogous to a two-deme island model in which 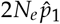 and 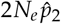 are the sizes of the two demes, and *M*_12_ and *M*_21_ are the scaled migration rates (e.g., Hudson and Kaplan 1988; Slatkin 1991; Nordborg 1997). Hereafter, we refer to the sub-population consisting of alleles associated with *A*_1_ or *A*_2_ as allelic class 1 or 2, respectively.

We can analyse this model efficiently using time-homogeneous phase-type theory (Hobolth *et al*. 2019). To this end, we define an intensity (rate) matrix as:

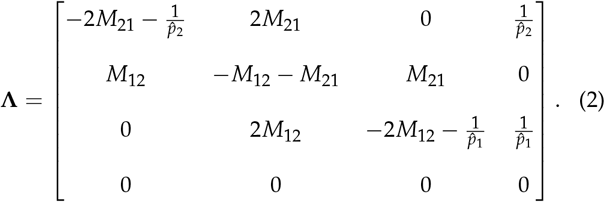

The first three rows in **Λ** are for states 1, 2, and 3, respectively. In row *i* (*i* ∈ {1,2,3}), the *j*–th element is the rate of jumping from state *i* to state *j* (*j ≠ i* and *j* ∈ {1,2,3,4}), and the diagonal element is the negative of the sum of all the other elements in this row. All elements of the last row of **Λ** are zero because state 4 is absorbing, so that the rate of leaving it is zero.

We can write **Λ** in a more compact form:

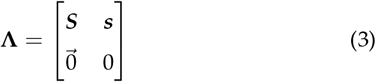

where ***S*** represents the 3-by-3 sub-matrix in the upper left corner of **Λ**, 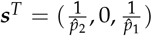 consists of the first three elements in the last column of **Λ** (the superscript *T* denotes matrix transposition), and 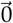 is a row vector of zeros. Thus, ***S*** contains the transition rates between the transient states, and ***s*** contains the rates of jumping to the absorbing state. ***S*** and ***s*** are referred to as the sub-intensity matrix and the exit rate vector, respectively.

Assume that *i* and 2 – *i* alleles in the sample are associated with *A*_1_ and *A*_2_, respectively. The time it takes for the process to reach the most recent common ancestor (MRCA) of the pair of alleles is a random variable that follows a phase-type distribution (Bladt and Nielsen 2017; Hobolth *et al*. 2019). To calculate the expected value of this random variable, denoted by *T*_*i*,2−*i*_, we define the Green’s matrix ***U*** = {*u_ij_*}, where *u_ij_* is the expected amount of time the process spends in state *j* prior to reaching the MRCA, provided that the initial state is *i* (*i, j* ∈ {1,2,3}). As shown in Supplementary Text S.2, ***U*** can be calculated as:

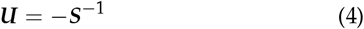

(see also Theorem 3.1.14 in Bladt and Nielsen (2017)). Take *T*_0,2_ as an example. The sample is in state 1. The expected amount of time the coalescent process spends in state *k* before reaching the MRCA is *u*_1*k*_ (*k* ∈ {1,2,3}). Thus, 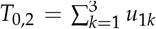. More generally, we have

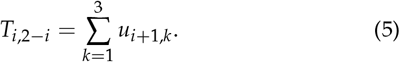

It is also possible to use phase-type theory to obtain the probability density function and all the moments of the coalescence time (Hobolth *et al*. 2019).

Define the initial condition vector as ***α*** = (*α*_1_, *α*_2_, *α*_3_), where *α_i_* is the probability that the sample is in state 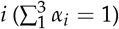. Thus, for *T*_0,2_, ***α*** = (1,0,0). Further let ***D**^T^* = (1,1,1). We can rewrite (5) as:

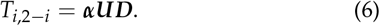

As we will see later, expressing the results this way allows us to accommodate non-equilibrium situations. The vector ***D*** is known as the reward vector. Its *k*-th element *D_k_* is the reward rate for every unit time the process stays in state *k*, such that the total contribution to *T*_*i*,2–*i*_ made by state *k* is *u*_*i*+1,*k*_*D_k_*.

It is possible to obtain ***U*** analytically for the general model with reversible mutation between *A*_1_ and **A**_2_, as specified by (2). However, its terms are complicated, and are not shown. For sites that are not very tightly linked to the selected locus, movements of lineages between the two allelic classes are primarily driven by recombination (i.e., *ρ* ≫ *μ_ij_*). Furthermore, with only two alleles at the selected locus, the general model is most appropriate for cases where the selected locus contains a small handful of nucleotides. In this case *μ_ij_* is of the order of the average nucleotide diversity at neutral sites (e.g., about 0.02 in *Drosophila melanogaster* or about 0.001 in humans). For most applications, therefore, it is sufficient to work with a simplified model with *μ_ij_* = 0. In this case, we have 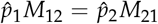 (i.e., there is conservative migration; Nagylaki (1980)), which leads to:

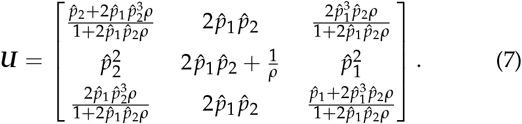

Summing the three rows, we have:

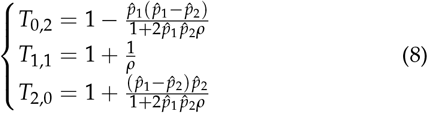

The results in (8) are the same as those derived by Nordborg (1997). The additional insight obtained here is given by (7). For instance, regardless of whether the initial state is 1 or 3, the process spends, on average, an equal amount of time in state 2 before coalescence (i.e., *u*_12_ = *u*_32_ in (7)). The results presented in Figure S1 further confirm that the simplified model should suffice in most cases, because the general model is well approximated by the simplified model for large enough *ρ*.

Let *π*_*i*,2−*i*_ be the expected diversity when *i* and 2 − *i* alleles in the sample are associated with *A*_1_ and *A*_2_, respectively. Under the infinite sites model (Kimura 1969), *π*_*i*,2−*i*_ = 2*θT*_*i*,2−*i*_, where *θ* = 2*N_e_v* and *v* is the mutation rate per generation at the neutral site. To put the discussion in context, we note that the expected coalescence time for two alleles is 1 under the neutral model with constant population size. From (8), we can see that *T*_1,1_ is independent of 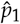 and 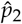, and is always greater than 1. For *T*_0,2_, it is < 1 or > 1 when 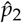 is < 0.5 or > 0.5, respectively. Similarly, *T*_2,0_ is < 1 or > 1 when 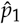 is < 0.5 or > 0.5, respectively. These trends hold even when there is reversible mutation between *A*_1_ and *A*_2_ (Figure S1).

In reality, the selected variants are often unknown, and detecting targets of balancing selection typically relies on investigating how diversity levels change along the chromosome (Charlesworth 2006; Fijarczyk and Babik 2015). It is therefore useful to consider the expected coalescence time for two randomly sampled alleles at the neutral site, defined as:

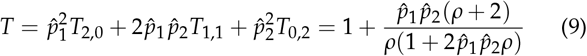

where the results in (8) are used. The nucleotide site diversity is given by *π* = 2*Tθ*. Figure 2 shows that the diversity level is highest when 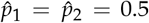. This is also true when there is reversible mutation between *A*_1_ and *A*_2_ (Figure S2). The simplified model is inherently symmetrical. For example, the curve for 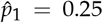 is identical to that for 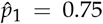. In all cases, marked effects on diversity are only seen in the immediate genomic neighbourhood of the selected site where *ρ* is of order 1 or less.

**Figure 2.**
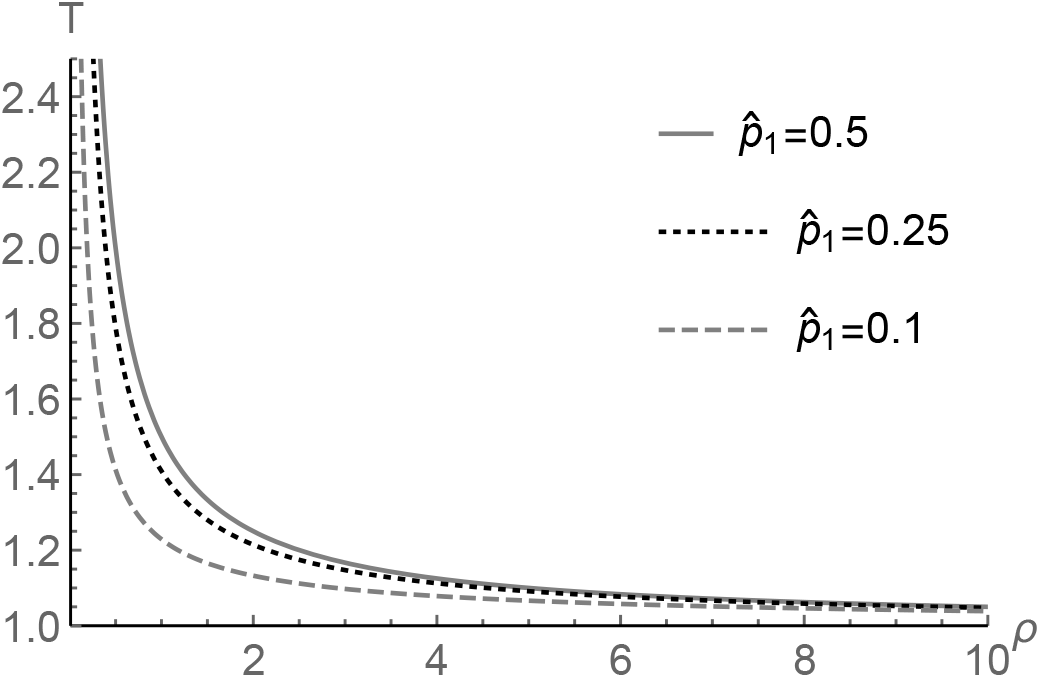
The expected pairwise coalescence time as a function of *ρ*. The simplified model with *μ*_12_ = *μ*_21_ = 0 is considered. 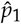 is the equilibrium frequency of *A*_1_at the selected locus.

### LD between the selected locus and a linked neutral site

The expected pairwise coalescence time obtained in the previous section can be used to calculate a measure of LD between the two loci (Charlesworth *et al*. 1997). Assume that the neutral locus is segregating for two variants *B*_1_ and *B*_2_. Let the frequencies of *B*_1_ in allelic class 1 and 2 be *x* and *y*, respectively. Thus, the frequency of *B*_1_ in the population is 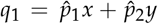, and that of *B*_2_ is *q*_2_ = 1 – *q*_1_. Let *δ = x – y*. The coefficient of LD between the two loci is given by 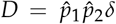 (see p. 410 of Charlesworth and Charlesworth 2010). The corresponding correlation coefficient is 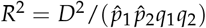. It is impossible to derive a simple expression for 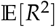. A widely-used alternative can be written as:

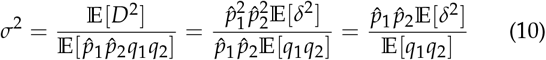

where we have used the fact that 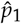 and 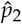 are assumed to be constant (Ohta and Kimura 1971; Strobeck 1983; McVean 2002). Note that 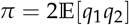 is the expected diversity at the neutral site.

As discussed in the previous section, we have *π* = 2*θT* under the infinite sites model. To relate *E*[*δ*^2^] to the expected pairwise coalescence times, we first define the expected diversity within allelic class 1 and allelic class 2 as 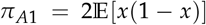 and 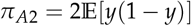, respectively. Again, under the infinite sites model, we have *π*_*A*1_ = 2*θT*_2,0_ and *π*_*A*2_ = 2*θT*_0,2_. In addition, let the weighted within allelic class diversity be 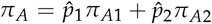. Note that 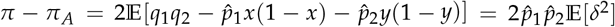. Inserting these results into the right-most term of (10), we have:

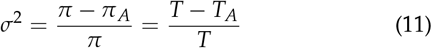

where 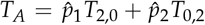 is the weighted average within allelic class coalescence time. Note that *σ*^2^ has the same form as the fixation indices (e.g., *F_ST_*) widely used in studies of structured populations. This close relationship between LD and the fixation indices was first pointed out by Charlesworth *et al*. (1997), who referred to *σ*^2^ as *F_AT_*. Our treatment here clarifies the relevant statements in this previous study. It also provides a genealogical interpretation of the results of Strobeck (1983).

Figure 3 shows *σ*^2^ as a function of *ρ* generated under the simplified model with *μ*_12_ = *μ*_21_ = 0. The level of LD between the selected and neutral loci is highest when 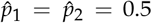, and decreases as 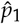 moves close to either 0 or 1 (note that the model is symmetrical such that, for 0 < *z* < 1, the curve for 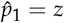 is identical to that for 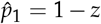). As expected, reversible mutation between *A*_1_ and *A*_2_ lowers LD by increasing the rate at which lineages move between the two allelic classes (Figure S3). These results mirror those described above for diversity levels. Together they show that the effect of balancing selection on linked diversity and LD patterns is largest when the equilibrium frequencies of the selected variants are close to 50%.

**Figure 3.**
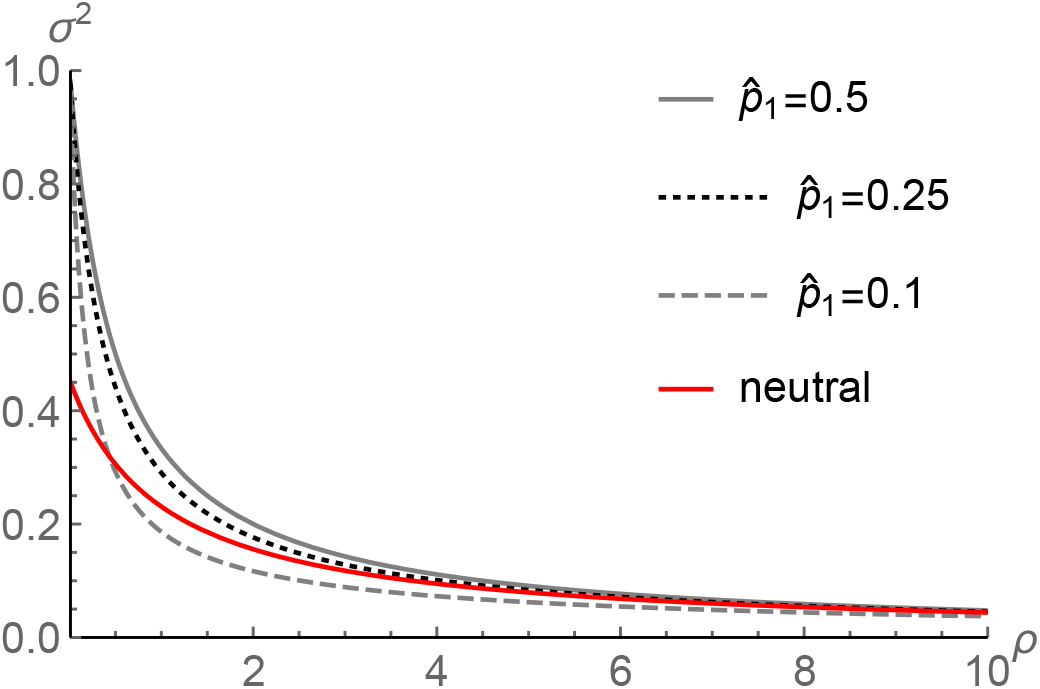
The level of LD between the selected and neutral loci as a function of *ρ*. The simplified model with *μ*_12_ = *μ*_21_ = 0 is considered. The neutral expectation for *σ*^2^ is also included.

It is informative to compare LD patterns under balancing selection with those under neutrality (i.e., *σ*^2^ = (5 + *ρ*)/(11 + 13*ρ* + 2*ρ*^2^); Ohta and Kimura 1971). With balancing selection and 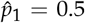, elevated LD is observed when *ρ* < 4 (Figure 3). With 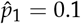, LD is higher than neutral expectation when *ρ* < 0.5, and it becomes lower than the neutral level when *ρ* > 0.5. Considering crossing over alone, the scaled recombination rate per site is of the order of 0.002 in humans, and 0.01 in *Drosophila*.These values go up substantially if we also take into account gene conversion (e.g., Campos and Charlesworth 2019). Thus, even when the effect of balancing selection is at its maximum, the region affected is small. The effect becomes rather insubstantial when the equilibrium frequency is close to 0 or 1, suggesting that such selection targets are probably extremely difficult to detect.

### Total branch length

We now consider the situation when a sample of *n* alleles is available, with *n*_1_ of them associated with *A*_1_ and *n*_2_ with *A*_2_(*n*_1_ + *n*_2_ = *n*). Let *L*_*n*_1_,*n*_2__ be the expected total branch length of the gene tree that describes the ancestry of the sample with respect to a neutral site linked to the selected locus. Note that, when *n* = 2, *L*_*n*_1_,*n*_2__ = 2*T*_*n*_1_,*n*_2__. The results in Figure 2 imply that close genetic linkage to a locus under balancing selection will result in an increase in the total branch length. Because the expected number of segregating sites in the sample is given by *θL*_*n*_1_,*n*_2__ under the infinite sites model, we expect to see more polymorphic sites in regions surrounding targets of long-term balancing selection. This theoretical expectation underlies several tests for balancing selection (Hudson *et al*. 1987; DeGiorgio *et al*. 2014).

To illustrate the calculation, consider a sample with three alleles. It can be in one of four possible states, with states 1, 2, 3, and 4 corresponding to situations where 0, 1, 2, and 3 of the sampled alleles are associated with *A*_1_. Going backwards in time, the coalescent process can move between these states via recombination or mutation between allelic classes. For instance, in state 1 all three alleles are associated with *A*_2_, and the process moves to state 2 at rate 3*M*_21_. When more than one allele is in the same allelic class, coalescence may occur. Again, take state 1 as an example. There are three alleles in allelic class 2, so that the rate of coalescence is 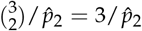. A coalescent event reduces the number of alleles to two, and thus moves the process to one of the three transient states depicted in Figure 1, referred to as states 5, 6, and 7 here. The transition rates between these states, as well as the rates of entering the absorbing state (i.e., the MRCA), are identical to those discussed above (i.e., (2)).

A diagram showing the transition rates between the states in this model can be found in Figure S4. The intensity matrix **Λ** for this model can be defined in the same way as described above, and is displayed in Supplementary Text S.3. **Λ** has a block structure:

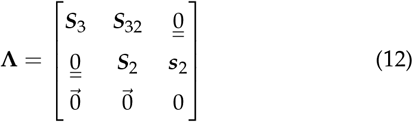

where 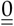 is a matrix of zeros. ***S***_3_ is a 4-by-4 matrix and contains the transition rates between states 1 - 4, all with three alleles. ***S***_32_ is a 4-by-3 matrix and contains the rates of coalescent events that move the process from a state with three alleles to one with only two alleles (i.e., from states 1 - 4 to states 5 - 7). Finally, ***S***_2_ and ***s***_2_ are the same as the corresponding elements defined in (3). The sub-intensity matrix ***S*** is the 7-by-7 sub-matrix in the upper left corner of **Λ**, and contains the transition rates between all the transient states.

Taking advantage of the block structure, we can calculate the Green’s matrix more efficiently as:

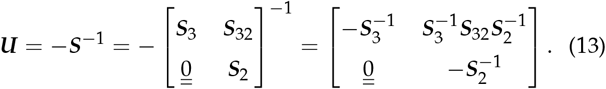

Recall that ***U*** = {*u_ij_*} and *u_ij_* is the expected amount of time the process spends in (transient) state *j* prior to reaching the MRCA, provided that the initial state is *i*. If, for instance, we want to calculate *L*_0,3_, we first note that the sample is in state 1. The process spends, on average, 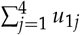 in states 1 - 4. Because these states have three alleles, the coalescent genealogy must have three lineages. Thus, these four states contribute 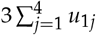 to *L*_0,3_. Similarly, states 5 - 7, which contain two alleles, contribute 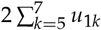. Putting these together, we have:

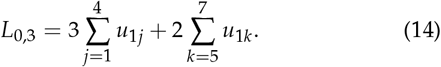

More generally, if the sample is in state *i*, we can define the initial condition vector as ***α*** = *e_i_*, where *i* ∈{1,2,3,4}and *e_i_* is a 1-by-7 vector whose elements are 0 except that the *i*-th element is 1. If we further define the reward vector as ***D***^*T*^ = (3, 3, 3, 3, 2, 2, 2), we have:

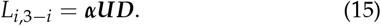

Note that this has the same form as (6). It is also possible to use phase-type theory to obtain the distribution and all the moments of the total branch length (Hobolth *et al*. 2019).

The approach can be easily extended to an arbitrary sample size *n*. As discussed above (see (9)), for data analysis, it is useful to consider the expected total branch length for a random sample of *n* alleles, defined as:

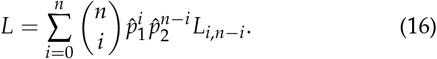

In Figure 4, we display *L* for several combinations of sample sizes and variant frequencies at the selected locus. To make the diversity-elevating effect more visible, we divide *L* by its neutral expectation (i.e., 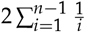). It is evident that, as *n* becomes larger, the sensitivity of *L* to 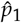 decreases, to the extent that, when *n* = 30, *L* is effectively independent of 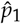. In addition, the strongest signal of elevated diversity appears when *n* = 2 and 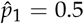, but becomes less pronounced as *n* increases. To interpret these observations, recall that, when *n* = 2, *π* = *θL*, whereas for larger *n, θL* is the expected number of segregating sites in the sample, denoted by 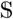. In data analysis, the nucleotide site diversity *π* is typically estimated from samples containing many alleles, and is known to be most sensitive to intermediate frequency variants (Tajima 1989). On the other hand, 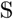 is determined primarily by low frequency variants in the sample. Thus, these results suggest that 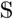 is less informative about balancing selection than *π*. However, the contrast between 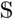 and *π* can be used as an index of the departure of the SFS from its expectation at neutral equilibrium (Tajima 1989). This clearly points to the importance of considering the SFS, which is done in the next subsection.

**Figure 4.**
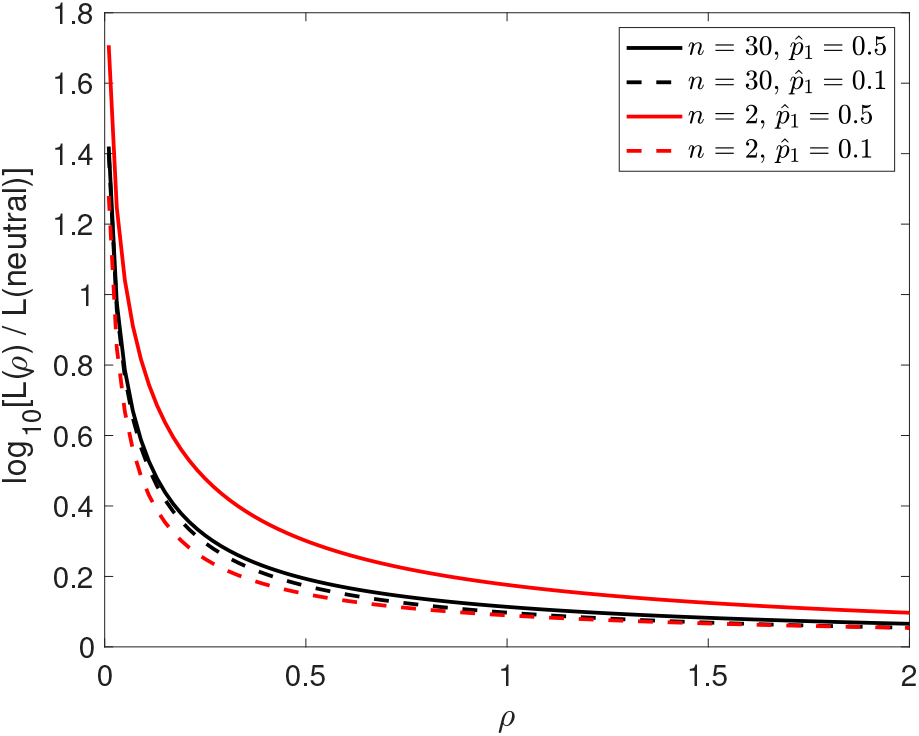
The expected total branch length *L* for several combinations of sample size (*n*) and equilibrium frequency of the selected variant 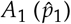. The value of *L* under balancing selection is divided by its neutral expectation. The y-axis is on the log_10_ scale.

This way of obtaining the total branch length is an alternative to the recursion method used in previous studies (Hudson and Kaplan 1988; DeGiorgio *et al*. 2014). The advantage of the current approach is that it can be extended to accommodate non-equilibrium dynamics such as population size changes and recent selection (see below). The dimension of the sub-intensity matrix ***S*** is 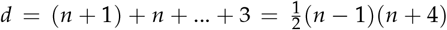. The numerical complexity increases rapidly because numerical matrix inversion requires *O*(*n*^6^) operations. However, by making use of the block structure (e.g., (13)), the number of operations is reduced to *O*(*n*^5^). Thus, this approach is computationally feasible for samples of dozens of alleles.

### The site frequency spectrum (SFS)

Again, consider a sample of *n* alleles at the neutral site, with *n*_1_ and *n*_2_ of them associated with *A*_1_ and *A*_2_, respectively. The *i*-th element of the SFS is defined as the expected number of segregating sites where the derived variant appears *i* times in the sample (0 < *i* < *n*). Note that this definition is different from the standard definition for a panmictic population in that it is conditional on *n*_1_ and *n*_2_. Consider the gene tree for the sample. We refer to a lineage (branch) that is ancestral to *i* alleles in the sample as a lineage of size *i* (0 < *i* < *n*). Under the infinite sites model, mutations on a lineage of size *i* segregate at frequency *i* in the sample. Let *φ_i_*(*n*_1_, *n*_2_) be the expected total length of all lineages of size *i* in the gene tree. The SFS under the infinite sites model can be expressed as *X_i_*(*n*_1_, *n*_2_) = *θφ_i_*(*n*_1_, *n*_2_) (e.g., Polanski and Kimmel 2003). We can calculate *ϕ_i_*(*n*_1_, *n*_2_) using phase-type theory with additional book keeping.

To illustrate the calculation, consider a sample of three alleles. Going backwards in time, before the first coalescent event, all the lineages are size one. After the first coalescent event, one lineage is size two, and the other is size one. Thus, the transient states of the coalescent process can be represented by 4-tuples of the form (*a*_1,1_, *a*_1,2_, *a*_2,1_, *a*_2,2_) where *a_i,j_* is the number of lineages of size *j* that are currently associated with *A_i_*. We have listed all the transient states in Table 1. The first four states contain three lineages, and the last four contain two lineages. We can determine the transition rates between the states using the same arguments that lead to Figures 1 and S4; the intensity matrix **Λ** is displayed in Supplementary Text S.4. Note that **Λ** has the same form as (12), so that we can obtain ***U*** using (13).

**Table 1.**
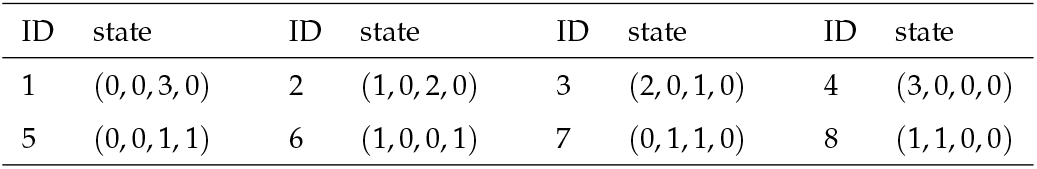
The transient states for a sample size of three

As an example, if *n*_1_ = 2 and *n*_2_ = 1, the starting state is 3, so that only the elements in the third row of ***U*** are relevant. Because states 1 - 4 contain three size one lineages, they contribute 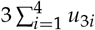 to *ϕ*_1_(2,1), but nothing to *φ*_2_(2,1). The last four states contain one size one lineage and one size two lineage. Thus, they contribute 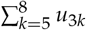 to both *φ*_1_(2,1) and *φ*_2_(2,1). Putting these results together, we have:

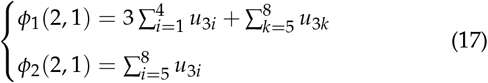

Define the initial condition vector ***α*** = (0,0,1,0,0,0,0,0), ***ϕ***(2,1) = (*ϕ*_1_ (2,1), *ϕ*_2_(2,1)) and

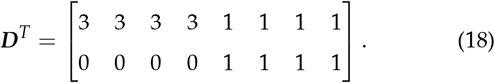

We have 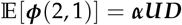, which is again in the same form as (6).

We can obtain the other ***ϕ***(*i*,3 – *i*) by defining the appropriate ***α***. In addition to the mean, it is also possible to use phase-type theory to obtain the variance of the SFS, as well as the covariance between different elements of the SFS (Hobolth *et al*. 2019). These results are applicable to any sample size *n* ≥ 2. We defer showing results regarding the SFS until a later section where a model of recent balancing selection is analysed.

Obtaining the SFS by working directly with the continuous time Markov process has been shown to be numerically more stable and accurate than approaches that rely on solving the diffusion equation numerically (Kern and Hey 2017). However, a limitation is that the size of the state space increases rapidly with *n*(Andersen *et al*. 2014). This is true even after exploiting the block structure of the sub-intensity matrix ***S***. For instance, when *n =* 16, the dimension of the largest sub-matrix in ***S*** is 922, but it increases to 3493 when *n =* 20. However, the flexibility of phase-type theory, especially its ability to accommodate complex non-equilibrium models, makes it a useful tool, as we show next.

## A model with strong balancing selection and changes in population size

So far we have only considered a model of balancing selection at statistical equilibrium. In this section, we switch our attention to a non-equilibrium model in which the population size changes in a stepwise manner. Specifically, we consider a diploid, randomly mating population. Looking back in time, its evolutionary history consists of *H* non-overlapping epochs, such that the effective population size is *N_e,h_* in epoch *h* (*h* ∈{1,2,…, *H*}). The duration of epoch *h* is [*t*_*h*–1_, *t_h_*), where *t*_0_ = 0 (the present) and *t_H_* = ∞. Thus, epoch *H*, the most ancestral epoch, has an infinite time span, over which the population is at statistical equilibrium. We assume that an autosomal locus is under balancing selection in epoch *H*, with two alleles *A*_1_ and *A*_2_ at equilibrium frequencies 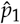 and 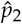, respectively. Based on the results shown in the previous sections, we only consider the simplified model without reversible mutation between *A*_1_ and *A*_2_. In addition, we assume that selection is sufficiently strong, and the changes in population size are sufficiently small, that the frequencies of the two alleles remain at 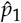 and 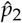 in the more recent epochs. A similar approach has been applied successfully to modelling the joint effects of background selection and demographic changes (Zeng 2013; Nicolaisen and Desai 2013; Zeng and Corcoran 2015).

### Total branch length

As before, consider a neutral site linked to the selected locus, with a sample of *n* alleles, of which *n*_1_ and *n*_2_ are associated with *A*_1_ and *A*_2_, respectively. Consider the expected total branch length, *L*_*n*1,*n*2_. Here time is scaled in units of 2*N*_*e*,1_ generations (twice the effective population size in the current epoch). We first note that the current model has the same states as the equilibrium model analysed above (e.g., see Figure S4 for *n* =3). The main difference between the two models lies in the transition rates between states.

We define the scaled recombination rate as *ρ* = 2*N*_*e*1_*r*. The rate at which an allele in allelic class *i* moves to allelic class *j* is 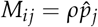. These have the same form as above (cf. Figure 1). In epoch *h*, the total number of alleles associated with *A*_1_ in the population is 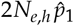. The probability that two alleles associated with *A*_1_ in the current generation coalesce in the previous generation is 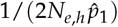. In other words, the probability that they remain un-coalesced for *z* generations is:

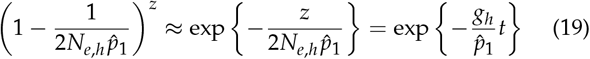

where *g_h_* = *N*_*e*,1_/*N*_*e,h*_ and *t* = *z*/(2*N*_*e*,1_). Thus, the coalescent rate between a pair of alleles in allelic class 1 is 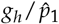 in epoch *h*. Similarly, the rate for two alleles in allelic class 2 is 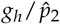.

In epoch *h*, the transition rates between the states are constant, and we can define an associated sub-intensity matrix, ***S***_*h*_. We have already noted that the states in the current model are the same as those in the equilibrium model. ***S***_*h*_ is very similar to the sub-intensity matrix for the equilibrium model (e.g., (12); see also Supplementary Text S.3). The only differences are (1) *ρ* is now defined as 2*N*_*e*1_*r* and (2) terms involving 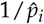 should be replaced by 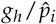.

Overall, the model has the following parameters: 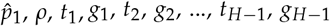, and *g_H_*. Among these, 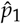 and *ρ* are shared across all the epochs, whereas epoch *h* has two epoch-specific parameters *t_h_* and *g_h_* (note that *t_H_* =∞). We have *H*sub-intensity matrices: ***S***_1_, ***S***_2_,…, ***S***_H_. In Supplementary Text S.5, we introduce time-inhomogeneous phase-type theory and prove the following result:

#### Theorem 1.

*Consider a continuous time Markov chain with finite state space* {1,2,…, *K, K*+1}, *where states* 1,…, *K are transient, and state K*+1 *is absorbing. Assume that the time interval* [0, ∞) *is subdivided into H non-overlapping epochs. The duration of epoch h is* [*t*_*h*–1_, *t_h_*), *where* 1 ≤*h*≤*H*, *t*_0_=0, *and t*_H_= ∞. *The sub-intensity matrix for epoch h is denoted by* ***S***_h_. *Then the Green’s matrix is*:

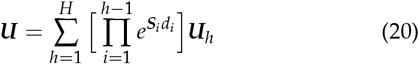

*where d_h_* = *t_h_* – *t*_*h*–1_, 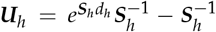 *and e^S_h_d_h_^* = 0 *if d_h_* = ∞.

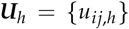 in (20) is the Green’s matrix for epoch *h*. Its element *u_ij,h_* is the expected amount of time the process stays in state *j* in this epoch if it enters the epoch in state *i*. Intuitively, *u_ij,h_* equals the amount of time the process spends in state *j* had the duration of epoch *h* been [0, ∞) (represented by 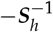) minus the amount of time it spends in state *j* in [*d_h_*, ∞) (represented by 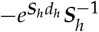). Let 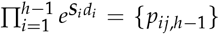. The element *p*_*ij,h*–1_ is the probability that the process starts from state *i* at *t*_0_=0 and is in state *j* by the end of epoch *h* - 1 at time *t*_*h*–1_. Thus, the overall Green’s matrix 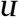 is the weighted mean of the epochs’ contributions, with the weights being the probabilities that the process enters the epochs in a particular state.

Applying this theorem requires the evaluation of matrix exponentials. Although this can be done analytically for certain models (e.g., Waltoft and Hobolth 2018), it is not feasible for the models considered here. We instead employ numerical methods (Al-Mohy and Higham 2010; Moler and Van Loan 2003), as implemented in the expm function in Matlab or the expm package in R. The computational cost for obtaining *e^***S***_*h*_*d_h_*^* is typically *O*(*d*^3^), where *d* is the dimension of ***S***_*h*_. Once ***U*** has been calculated, the expected total branch length is given by 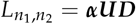 (see (15)).

In Figures 5a and b, we show *L*, the expected total branch length, for a random sample of *n* = 20 alleles (see (16)), under either a one-step population size increase or a one-step population size reduction. The population size change occurred at time *t* before the present. Because *L* is insensitive to 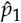 when *n* is relatively large (Figure 4), we only consider 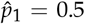 (the results are qualitatively very similar with *n* =2; not shown). Neutral diversity levels in genomic regions closely linked to the selected site are affected by recent population size changes to a much smaller extent than regions farther afield. This is because, for small ρ, migration of lineages between allelic classes is slow, such that the tree size is mainly determined by the divergence between allelic classes rather than drift within allelic classes. The importance of the divergence component increases with decreasing *ρ*. In particular, when there has been a recent reduction in population size, this effect protects against the loss of neutral polymorphisms in a larger genomic region (Figure 5b). Consequently, all else being equal, strong balancing selection affects a bigger stretch of the genome and produces a higher peak of diversity in smaller populations, potentially making them easier to detect. A similar observation has been made in models with self-fertilisation and background selection (Nordborg *et al*. 1996).

**Figure 5.**
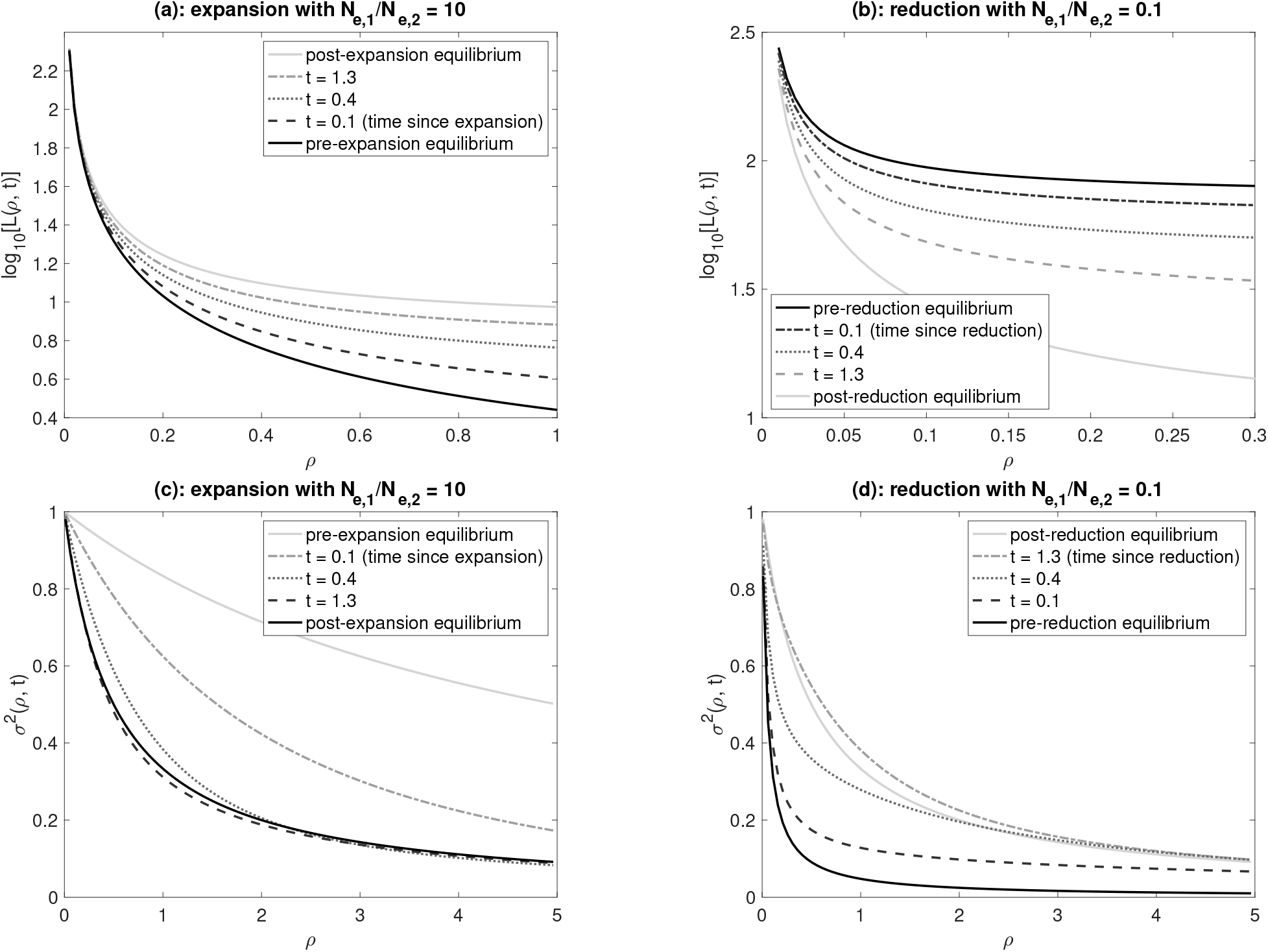
Expected total branch length and LD as a function of *ρ* and *t*. The population experienced a one-step change in population size at time *t* before the present. The population size in the present and ancestral epochs are *N*_*e*,1_ and *N*_*e*,2_, respectively. Time is scaled in units of 2*N*_*e*,1_ generations. The selected alleles *A*_1_ and *A*_2_ are at equilibrium frequencies 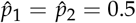. The sample size is *n* = 20.

Note that, although we have focused on calculating the total branch length, Theorem 1 can also be used to calculate the SFS. This can be done by defining an appropriate state space (e.g., Table 1) and a suitable reward matrix (e.g., (18)). We will demonstrate these calculations later when we analyse a model of recent balancing selection.

### LD between the selected locus and a linked neutral site

The measure of LD can be calculated by replacing *T* and *T_A_* in (11) with *T*(*t*) and *T_A_*(*t*). In Figures 5c and d, we can see that *σ*^2^ converges to its new equilibrium level at a much higher rate than the level of diversity, which is a well-known effect (e.g., McVean 2002). Interestingly, *σ*^2^ appears to approach its new equilibrium in a non-monotonic way. For instance, in Figure 5c, LD levels at *t* = 0.4 are temporarily higher than the equilibrium value (the solid black curve), but become lower than the equilibrium value at *t* = 1.3. In Figure 5d, we can see that the level of LD is higher, and extends further, after the population size reduction (see also Figure S5). These results further suggest that balancing selection may be easier to detect in smaller populations.

### Simulations

The theory developed above assumes that the frequencies of *A*_1_ and *A*_2_ remain constant at 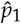 and 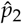, respectively. This is true only when the population size is infinite. With a finite population size, allele frequencies fluctuate around their equilibrium values due to genetic drift. To investigate the effects of stochastic allele frequency fluctuation on the accuracy of our model predictions, we conducted simulations using mbs (Teshima and Innan 2009). Briefly, each simulation replicate contained two steps: (1) forward simulation to obtain allele frequency trajectories for the selected variants given the demographic history; (2) coalescent simulation for a sample of *n* alleles at a linked neutral site, conditioning on the trajectories obtained in step 1 (see Supplementary Text S.6 for more details). Because the theory does not depend on a specific selection model, we used an overdominance model whereby the fitnesses of the three genotypes *A*_1_*A*_1_, *A*_1_*A*_2_, and *A*_2_*A*_2_ are 1 –*s*_1_, 1, and 1 –*s*_2_, respectively. The equilibrium frequencies are 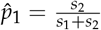>. and 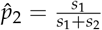.

To check the results presented in Figure 5, we let *s*_1_ = *s*_2_ = *s*, such that 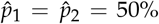. To simulate the population expansion model in 5a, we assumed that *N*_*e*,1_ = 20,000 (the effective population size of the current epoch) and *N*_*e*,2_ = 2,000 (the effective population size of the ancestral epoch). For the population reduction model in 5b, we used *N*_*e*,1_ = 2,000 and *N*_*e*,2_ = 20,000.

As shown in Figure S6, the theoretical predictions are highly accurate. Here selection was strong, as measured by *γ_min_* = 2*N*_*e,min*_*s* = 250, where *N*_*e,min*_ = min(*N*_*e*,1_, *N*_*e*,2_). To further check the robustness of our results, we reduced *s*, such that *γ_min_*= 20. The substantial reduction in the intensity of selection leads to a significantly higher level of fluctuation in the frequencies of the selected variants (Figure S7). Encouragingly, the theoretical predictions remain accurate (Table S1).

## A model of recent balanced polymorphism

We now turn our attention to the effects of the recent origin of a balanced polymorphism on patterns of genetic variability. Consider a diploid panmictic population with constant effective population size *N_e_*. At an autosomal locus, a mutation from *A*_1_(the wild type) to *A*_2_ (the mutant) arises. The fitnesses of the genotypes *A*_1_*A*_1_, *A*_1_*A*_2_, and *A*_2_*A*_2_ are *w*_11_ = 1 – *s*_1_, *w*_12_ = 1, and *w*_22_ = 1 – *s*_2_(*s*_1_ > 0 and *s*_2_ > 0; i.e., there is heterozygote advantage). As above, we ignore reversible mutation between *A*_1_ and *A*_2_. In what follows, we first use a forward-in-time approach to obtain equations for describing the increase in the frequency of *A*_2_ in the population. We then use the backwardin-time coalescent approach to calculate various measures of sequence variability in linked genomic regions. Wherever appropriate, we present results from a comparable selective sweep model, so that the two models can be compared.

### Frequency of the mutant allele in the population

Let the frequencies of *A*_1_ and *A*_2_ in the current generation be *p*_1_ and *p*_2_, respectively. Let 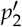 be the frequency of *A*_2_ in the next generation. Using the standard theory (reviewed in Chap. 2 of Charlesworth and Charlesworth (2010)), the change in allele frequency in one generation due to selection is given by

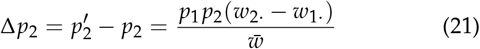

where *w*_1_ = *p*_1_*w*_11_ +*p*_2_*w*_12_, *w*_2_. = *p*_1_*w*_12_ + *p*_2_*w*_22_, and 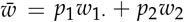. Assuming that both *s*_1_ ≪ 1 and *s*_2_ ≪ 1, Δ*p*_2_ ≈ *p*_1_*p*_2_(*w*_2_. – *w*_1_1) = *p*_1_*p*_2_(*p*_1_*s*_1_–*p*_2_*s*_2_). At equilibrium, Δ*p*_2_=0, such that the frequencies are 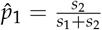 and 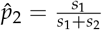.

When *p*_2_ ≪ 1, Δ*p*_2_ ≈ *s*_1_*p*_2_. This is the same as when *A*_2_ is under positive selection with fitnesses of the three genotypes being *w*_11_ = 1, *w*_12_ = 1 + *s*_1_, and *w*_22_ = 1 +2 *s*_1_, respectively (i.e., there is semi-dominance). Thus, we expect the initial signals generated by the increase in *p*_2_ to be similar to those from an incomplete selective sweep, referred to here as the “corresponding sweep model”.

The similarity between the two selection models means that we can borrow useful results from the selective sweep literature. In particular, after *A*_2_ has been generated by mutation, its frequency must increase rapidly for it to escape stochastic loss. Following an approach first proposed by Maynard Smith (1976), we assume that *p*_2_ increases instantly to 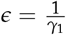, where *γ*_1_ = 2*N*_*e*s_1__ (see also Desai and Fisher 2007). Thereafter, *p*_2_ changes deterministically until its rate of change becomes very slow near the equilibrium point, when the coalescent process (considered in the next sub-section) is effectively the same as at equilibrium. Measuring time in units of 2*N_e_* generation, *p*_2_(*t*) satisfies:

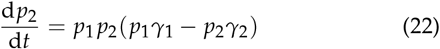

where *γ*_2_ = 2*N*_*e*_*s*_2_. The solution to this differential equation is

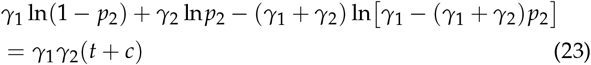

where *c* is a constant such that *p*_2_(0) = *ϵ*. We can obtain the frequency of *A*_2_ at time *t* by solving for *p*_2_ numerically.

It is instructive to compare the dynamics of *p*_2_(*t*) with those for the corresponding sweep model defined above. We assume that the frequency of the positively selected variant *A*_2_ increases instantly to *ϵ* and grows deterministically until 1 – *ϵ*. Let 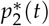 be the frequency of *A*_2_ at scaled time *t* after its frequency arrived at *ϵ*. It can be shown that:

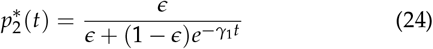

(Crow and Kimura 1970; Stephan *et al*. 1992).

A recent study explicitly considered the stochastic phases when the frequency of the positively selected variant *A*_2_ is below *ϵ* or greater than 1 _ *ϵ* (Charlesworth 2020a). These two phases contribute relatively little to the fixation time under the current model with strong selection and semi-dominance (see Table 1 of Charlesworth 2020a). Furthermore, when the frequency of *A*_2_ is very close to 0 or 1, the coalescent process is effectively the same as under neutrality. Thus, ignoring these two stochastic phases is reasonable for our purposes.

In Figure 6, we display three balancing selection models, all with *γ*_1_ = 500, but different *γ*_2_ values, so that they have different equilibrium allele frequencies. For comparison, the corresponding sweep model with *γ*_1_ = 500 is also presented. As can be seen, the allele frequency trajectories for the balancing selection models and the corresponding sweep model are similar only for a rather short period. After that, *p*_2_(*t*) increases at a much slower pace than 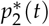. As shown below, these observations explain the differences between a recent balanced polymorphism and the spread of a beneficial mutation with respect to their effects on diversity patterns in nearby genomic regions.

**Figure 6.**
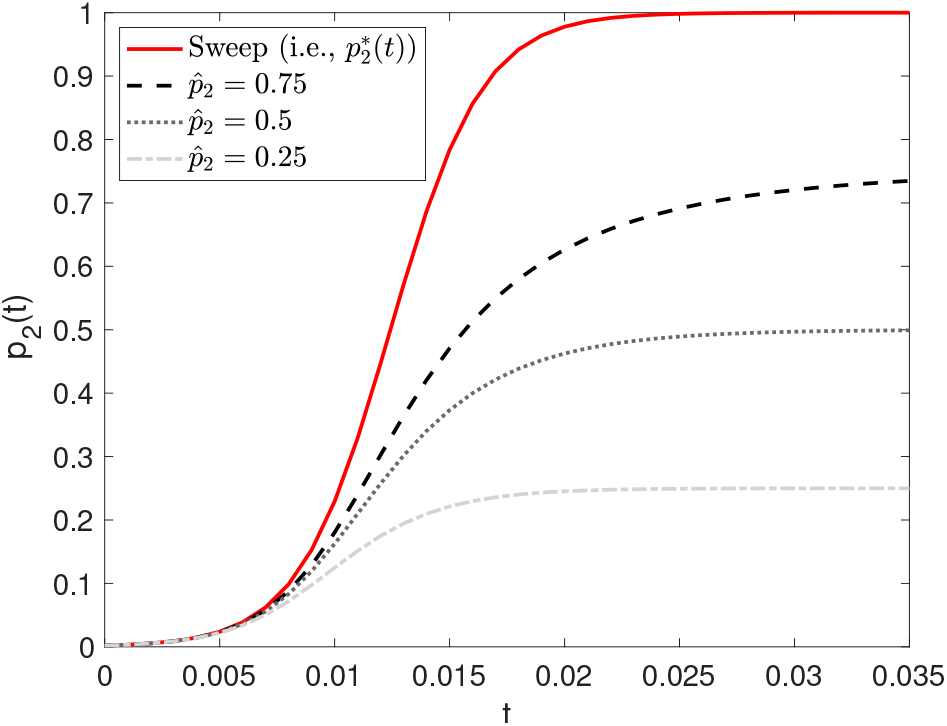
The frequency of the mutant allele *A*_2_ as a function of *t* (time since its frequency reached *ϵ*). *γ*_1_ = 500. *γ*_2_ is adjusted such that the equilibrium frequency 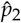 is 0.25, 0.5, and 0.75, respectively. The trajectory under the corresponding sweep model is included for comparison.

### Total branch length

We extend the coalescent approach developed above for the equilibrium model, in order to calculate the expected total branch length *L* for a random sample of *n* alleles at a linked neutral site (see (16)). The frequency of *A*_2_ at the time of sampling is *p*_2_(*t*) where *t* is the time since the frequency of *A*_2_ reached *ϵ*, expressed in units of 2*N_e_* generations. At time *τ* before the present (0 ≤*τ* < *t*), the frequency of *A*_2_is given by *p*_2_(*t* – *τ*). For *τ* ≥ *t*, the process reduces to a standard neutral coalescent model with constant population size. To make use of Theorem 1, we divide [*p*_2_(*t*), *ϵ*) into *H* – 1 equal-sized bins, such that the *h*-th bin is [*p*_2,*h*–1_, *p*_2,*h*_), where 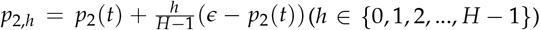. Let *τ_h_* be the solution to *p*_2_(*t* – *τ_h_*) = *p*_2*h*_ given by (23). The corresponding time interval for bin *h* is [*τ*_*h*–1_, *τ_h_*), which is shorter when the frequency of *A*_2_ is changing at a faster rate. Thus, we have *H* epochs, with the first *H* – 1 in [0, *t*) and epoch *H* covering the whole of [*t*, ∞) (Figure S8).

Consider epoch *h* with *h* < *H*. The state space in this epoch is the same as that discussed above for the equilibrium model (see the arguments leading to (12)). Thus, the sub-intensity matrix for this epoch, ***S***_*h*_, can be obtained in a similar way (cf., Figure S4). The only complication is that the frequency of *A*_2_ changes within the epoch. However, if the time interval is sufficiently small, we can treat the frequency of *A*_2_ as if it were constant. Here we set the frequency of *A*_2_ in epoch *h* to its harmonic mean *q*_2,*h*_, which can be calculated as:

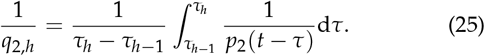

We can then obtain ***S***_*h*_ by simply replacing 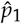 and 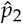 in the subintensity matrix for the equilibrium model with *q*_1,*h*_ and *q*_2,*h*_, where *q*_1,*h*_ = 1 – *q*_2,*h*_.

Note that, although the space state is the same for the epochs in [0, *t*), this is not true for the transition from epoch *H* – 1 to epoch H. At the end of epoch *H* – 1, if more than one allele is associated with *A*_2_, they coalesce into a single ancestral allele instantly. If the resulting ancestral allele is the only allele left, the process is terminated. Otherwise, if there are also *n*_1_ alleles associated with *A*_1_ at the time, then the *n*_1_ + 1 alleles enter epoch *H* and coalesce at rate 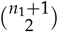. Thus, we need a mapping matrix *E*_*H*–1,*H*_, which is defined below (S22) in Supplementary Text S.5, to take into account the differences between the two epochs. For instance, for a sample of two alleles, the state space in [0, *t*) has three transient states: (0, 2), (1, 1), and (2, 0), where the two numbers of each tuple represent the number of alleles associated with *A*_1_ and *A*_2_, respectively. However, epoch *H* has only one transient state, representing two uncoalesced alleles. If the process is in state (0, 2) at the end of [0, *t*), it terminates with the instant coalescence of the two alleles. If the process is in any of the other two states, it enters epoch *H* with the same starting condition. Thus 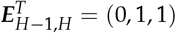, where 0 in the first element means it is impossible to enter epoch *H* via state 1 in epoch *H* – 1, and the 1s mean that, if the process is in state 2 or 3 by the end of epoch *H* – 1, the process begins epoch *H* in state 1.

In all, the model has the following parameters: *γ*_1_, *γ*_2_, *t*, and *ρ*. By increasing the number of bins in the discretisation scheme (i.e., *H*; Figure S8), we can get arbitrarily accurate approximations. The results presented below are based on values of *H* such that the size of the frequency bins is about 1%. This is a rather conservative choice; using larger bins does not significantly change the results. Once the sub-intensity matrices are defined (i.e., ***S***_*h*_ for 1 ≤ *h* ≤ *H*), we can obtain 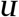 using Theorem 1 (see also Supplementary Text S.5) and 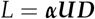 (see (15)).

Figure 7a shows how neutral diversity levels are affected by a recent balanced polymorphism, using the balancing selection model with 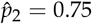 considered in Figure 6. Initially, the rapid increase in the frequency of *A*_2_ produces a drop in neutral diversity in nearby regions (the solid blue line). The maximum extent of reduction appears when *p*_2_(*t*) is close to its equilibrium value (the dotted line; *p*_2_(0.04) =0.742). After that, the diversity level starts to recover. Here, the increase in diversity level is fastest for regions closely linked to the selected site, because coalescence is slow when *ρ* is small. This leads to a U-shaped diversity pattern that persists for some time, which is followed by a rather slow approach to the equilibrium value (Figure S9). These dynamics are qualitatively the same when we consider a larger sample size with 20 alleles, although the reduction in diversity is less pronounced (Figure S10). Similar patterns are also observed for the other two balancing selection models in Figure 6 (Figure S11). The main difference is that models with a smaller 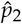 tend to result in a smaller reduction in neutral diversity. For instance, for the model with 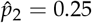, the maximum reduction in nucleotide site diversity in very tightly linked regions is less than 6% (as opposed to a more than 50% reduction in Figure 7a), potentially making them very difficult to detect from data.

**Figure 7.**
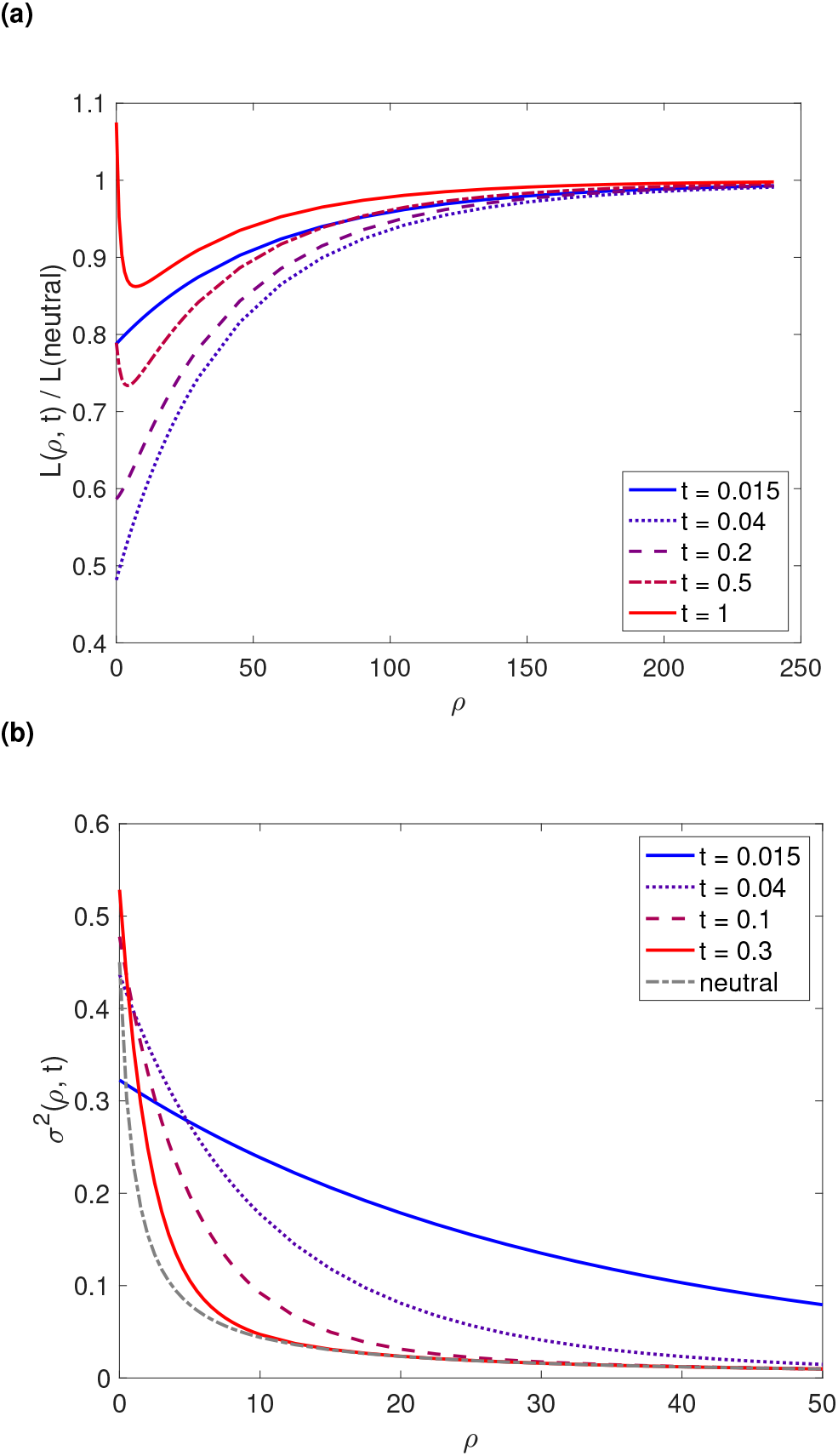
Nucleotide site diversity and LD in genomic regions surrounding a recently-emerged variant under balancing selection. The parameters are *γ*_1_ = 500 and 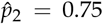 (as in Figure 6). The discretisation scheme has *H* = 76 bins. In (a), the expected total branch length for a sample of *n* = 2 alleles is calculated for various value of *t*, the time since the frequency of *A*_2_ reached *ϵ*. To make the effects more visible, *L* is divided by its neutral expectation. *σ*^2^ in (b) measures the level of LD between the selected locus and a linked neutral site. For comparison, the neutral expectation of *σ*^2^ is also included.

### LD between the selected locus and a linked neutral site

It is straightforward to use the method developed in the previous subsection to calculate *σ*^2^. From Figure 7b, we make two observations. First, LD builds up quickly and extends to a large genomic region when the frequency of *A*_2_ is increasing rapidly (blue solid curve vs the neutral curve). This suggests the formation of long haplotypes around the selected locus, which can be used to help detect selection targets, as is done in extended haplotype tests (e.g., Voight *et al*. 2006; Ferrer-Admetlla *et al*. 2014). Second, the level of LD starts to decline before the reduction in diversity is maximal (the dotted curves in Figures 7a and b), suggesting that LD based detection methods will have already lost a substantial amount of their statistical power by this time. This implies that LD and diversity patterns complement each other when it comes to detecting targets of recent balancing selection.

### Differences between balancing selection and selective sweeps in their effects on the total branch length and LD

We can analyse selective sweep models using the discretisation scheme outlined in Figure S8. In Figure 8a, we compare the balancing selection model shown in Figure 7 to its corresponding sweep model, with respect to their effects on *L* (the expected total branch length). Because the frequency of the beneficial allele increases much more rapidly (Figure 6), it causes a more pronounced reduction in diversity than the balanced polymorphism of the same age. Fixation of the beneficial allele occurs at *t* = 0.025. After that, diversity returns to its neutral level over a time period of the order of 2*N_e_* generations, which is much faster than the time it takes for diversity to reach its equilibrium level under balancing selection (Figure S9). The patterns are similar when a larger sample size is considered (Figure S12).

**Figure 8.**
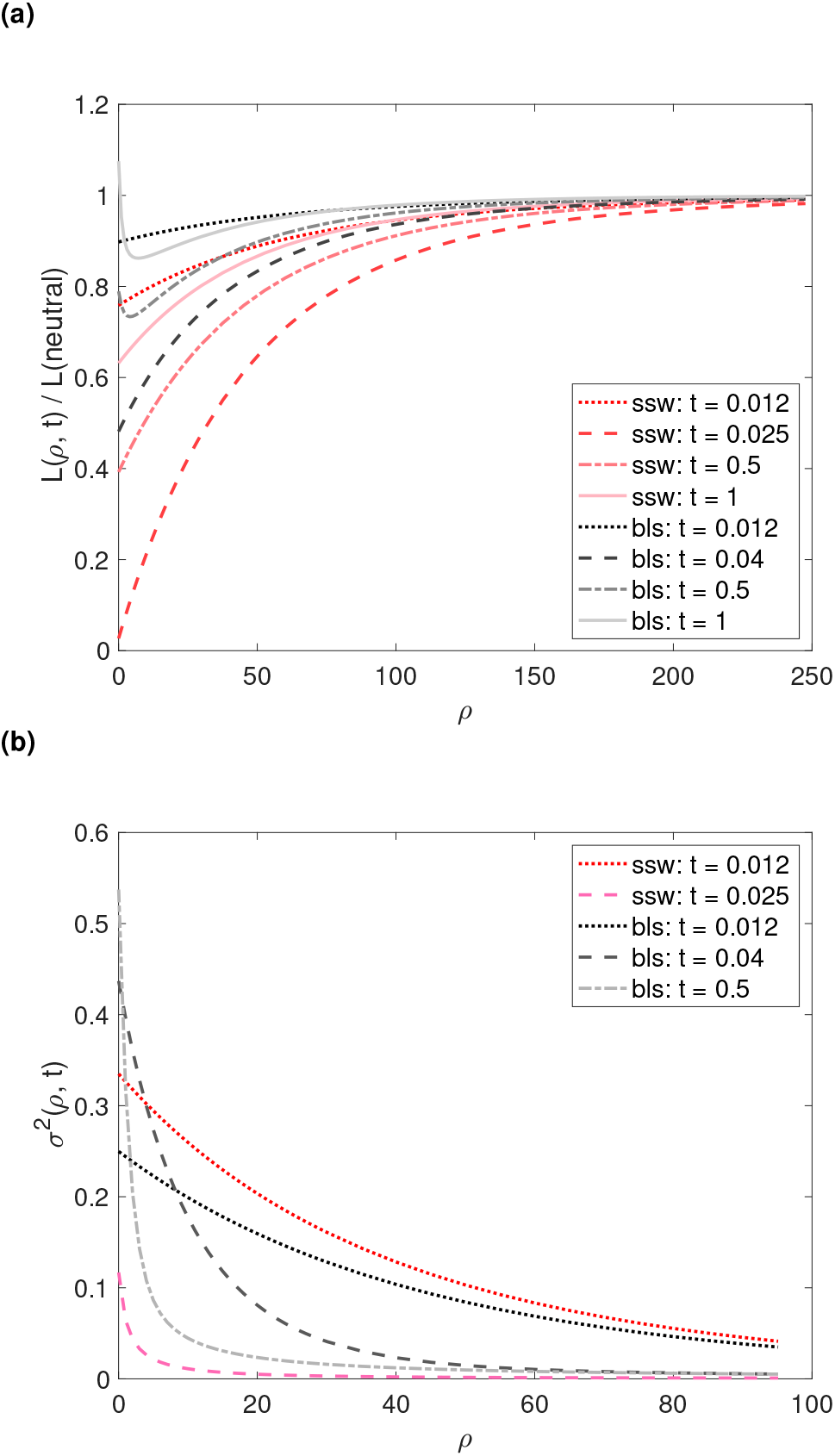
Comparing recent balancing selection with the corresponding sweep model, with respect to their effects on diversity and LD levels in surrounding genomic regions. The parameters of the balancing selection model (bls) are *γ*_1_ = 500 and 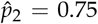 (i.e., the same as in Figure 7). The corresponding sweep model (ssw) has *γ*_1_ = 500. In (a), the expected total branch length for a sample of *n* = 2 alleles, divided by its neutral value, is presented. In (b), we consider the level of LD between the selected locus and a linked neutral site, as measured by *σ*^2^. Fixation (taken as the time when the mutant allele frequency reaches 1 – *ϵ*) occurs at *t* = 0.025 under the sweep model. The reduction in diversity reaches its maximum at *t* ≈ 0.04 under the balancing selection model.

A comparison between the two selection models with respect to their effects on LD patterns in the surrounding neutral region is shown in Figure 8b. Both models result in elevated LD. As expected, the corresponding sweep model leads to a more pronounced build-up of LD (red vs black dotted lines). This suggests that recent balancing selection is harder to detect than a comparable beneficial mutation. Under both models, LD starts to decay before the reduction in diversity is maximal (pink vs grey dashed lines). The decay appears to be much faster under the sweep model. This is because, under the balancing selection model, *A*_2_ approaches an equilibrium frequency, instead of fixation. Therefore, a sizeable genomic region remains at elevated levels of LD with the selected locus for a longer period. Recall that diversity levels also take much longer to reach equilibrium under balancing selection (Figure 8a). Thus, there may well be a bigger window of opportunity for detecting targets of recent balancing selection, despite the fact that the signals they produce tend to be less dramatic than those produced by the corresponding sweep model.

### The site frequency spectrum

The SFS can also be obtained using the time discretisation procedure. Here the state space is the same as that detailed for the equilibrium balancing selection model. As above, we obtain the sub-intensity matrix for epoch *h* by replacing 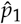 and 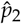 in the sub-intensity matrix for the equilibrium model (e.g., Supplementary Text S.4) with *q*_1,*h*_ and *q*_2,*h*_, respectively. We then use Theorem 1 to calculate *X_i_*(*n*_1_, *n*_2_). It is more instructive to consider the SFS for a sample of *n* randomly collected alleles, defined as:

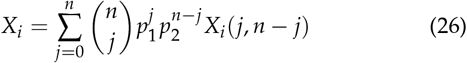

where *p*_1_ and *p*_2_ are the frequencies of *A*_1_ and *A*_2_ at the time of sampling. The effects selection has on the shape of the SFS are visualised using the ratio *X_i_/X_i_* (neutral), where *X_i_* (neutral) =2*θ*/*i*.

In Figure 9, we present the SFS at different time points since the arrival of the mutant allele, for both the balancing selection model and the corresponding sweep model considered in Figure 8. When the frequency of the selected variant is rapidly increasing in the population, both types of selection produce a U-shaped SFS, with an excess of both low and high frequency derived variants. The extent of distortion is maximised around the time when the reduction in neutral diversity is also the most pronounced (see plots in the second row). The corresponding sweep model has a much bigger effect on the shape of the SFS. For example, under the sweep model, at the time of fixation (*t* =0.025), *X_9_*/*X*_8_ = 4.91 and *X*_1_/*X*_2_ = 8.05. In contrast, when the SFS is most distorted under the balancing selection model (*t* =0.04), *X*_9_/*X*_8_ = 1.34 and *X*_1_/*X*_2_ = 3.29. The excess of high frequency derived variants quickly disappears after the selected allele has stopped its rapid increase in frequency (plots in the third row), although the SFS remains U-shaped for longer under balancing selection. The plots in the last row shows the transition from a situation with reduced diversity and an excess of low frequency variants to a situation that resembles the pattern expected under long-term balancing selection, with an elevated diversity level and an excess of intermediate frequency variants. Qualitatively similar dynamics have been observed for the balancing selection models with 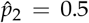 and 0.25, respectively (Figure S13). Again, the SFS-distorting effect is weaker when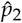 is smaller, with the case with 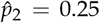 producing hardly any excess of low and high frequency variants even when *A*_2_ is increasing in frequency.

**Figure 9.**
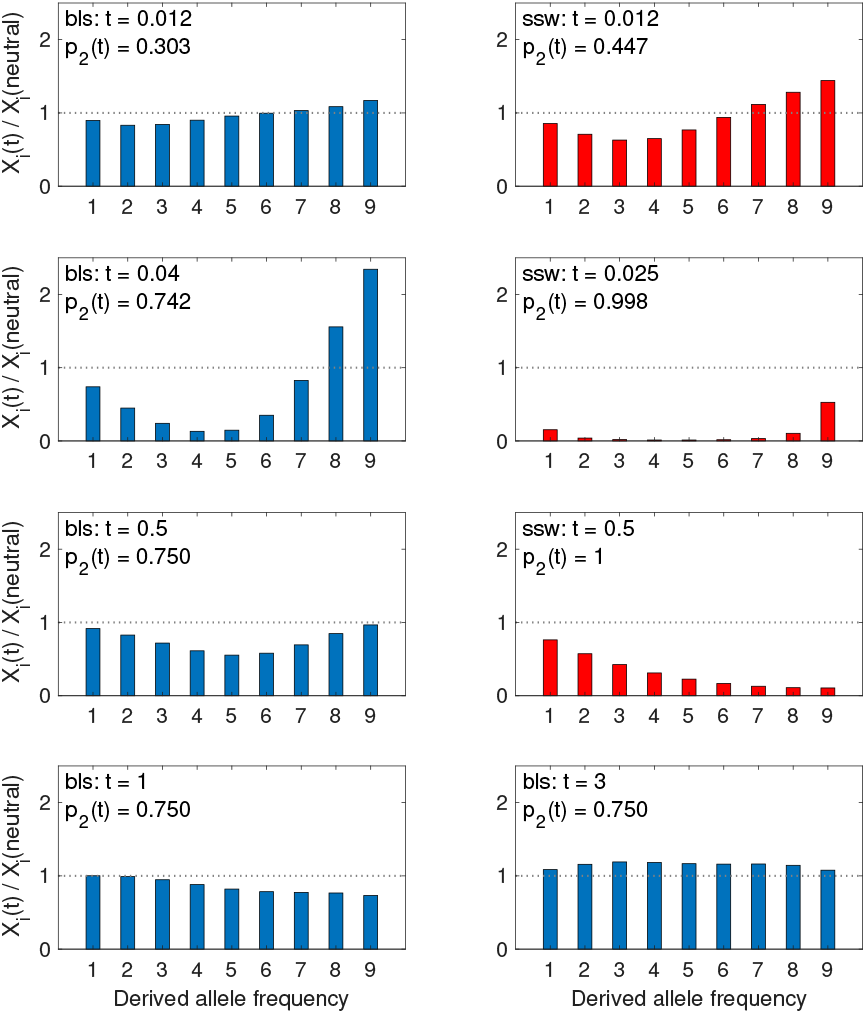
The SFS at various time points after the arrival of the selected variant for a random sample of 10 alleles. The balancing selection (bls) and selective sweep (ssw) models are the same as those shown in Figure 8. The scaled recombination frequency between the focal neutral site and the selected site is *ρ* = 2. The reduction in diversity reaches its maximum at *t* ≈ 0.04 and 0.025 (fixation) under the balancing selection and selective sweep models, respectively. The SFS under selection is expressed relative to its neutral expectation.

To investigate the SFS further, we consider *π* (the nucleotide site diversity) and Watterson’s *θ_W_*. Recall that, under the infinite sites model, *π* = 2*θT*, where *T* is defined by (9). Let 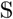 be the expected number of segregating sites in a sample of size *n*. We have 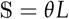. Because 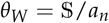 where 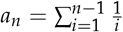, we have *θ*_*W*_ = *θL*/*a_n_*. Following Becher *et al*. (2020), we define

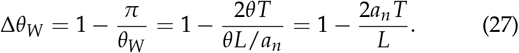

Δ*θ*_*w*_ = 0 under neutrality, >0 when there is an excess of rare variants, and < 0 when there is an excess of intermediate frequency variants.

Figure 10 shows Δ*θ*_*W*_ for the balancing selection model with *γ*_1_ = 500 and 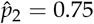 (as in Figures 6 - 9); the corresponding sweep model is also included for comparison. At *t* = 0.012, the balancing selection model produces no obvious deviation from neutrality (black dotted line), whereas the sweep model has already started to cause a significant excess of rare variants (red dotted line). This is consistent with the much slower increase in the frequency of *A*_2_ under balancing selection (*p*_2_(0.012) =0.303 vs 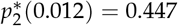). The extent of deviation caused by the sweep is maximal around the time when *A*_2_ becomes fixed (*t* ≈ 0.025; pink dashed line). Under the balancing selection model, the maximum deviation appears when the frequency of *A*_2_ becomes close to its equilibrium value (*t* ≈ 0.04; grey dashed line), but is less pronounced than under the sweep model. After the maximum is achieved, diversity patterns gradually return to neutrality over 4*N_e_* generations under the sweep model. For the balancing selection model, there is a much longer period of non-stationary dynamics as shown by the light blue and blue lines. The observations are qualitatively similar for the two sample sizes considered (*n* =10 vs *n* = 35). Nonetheless, the extent of deviation in the SFS is more conspicuous when *n* =35, suggesting an increase in statistical power.

**Figure 10.**
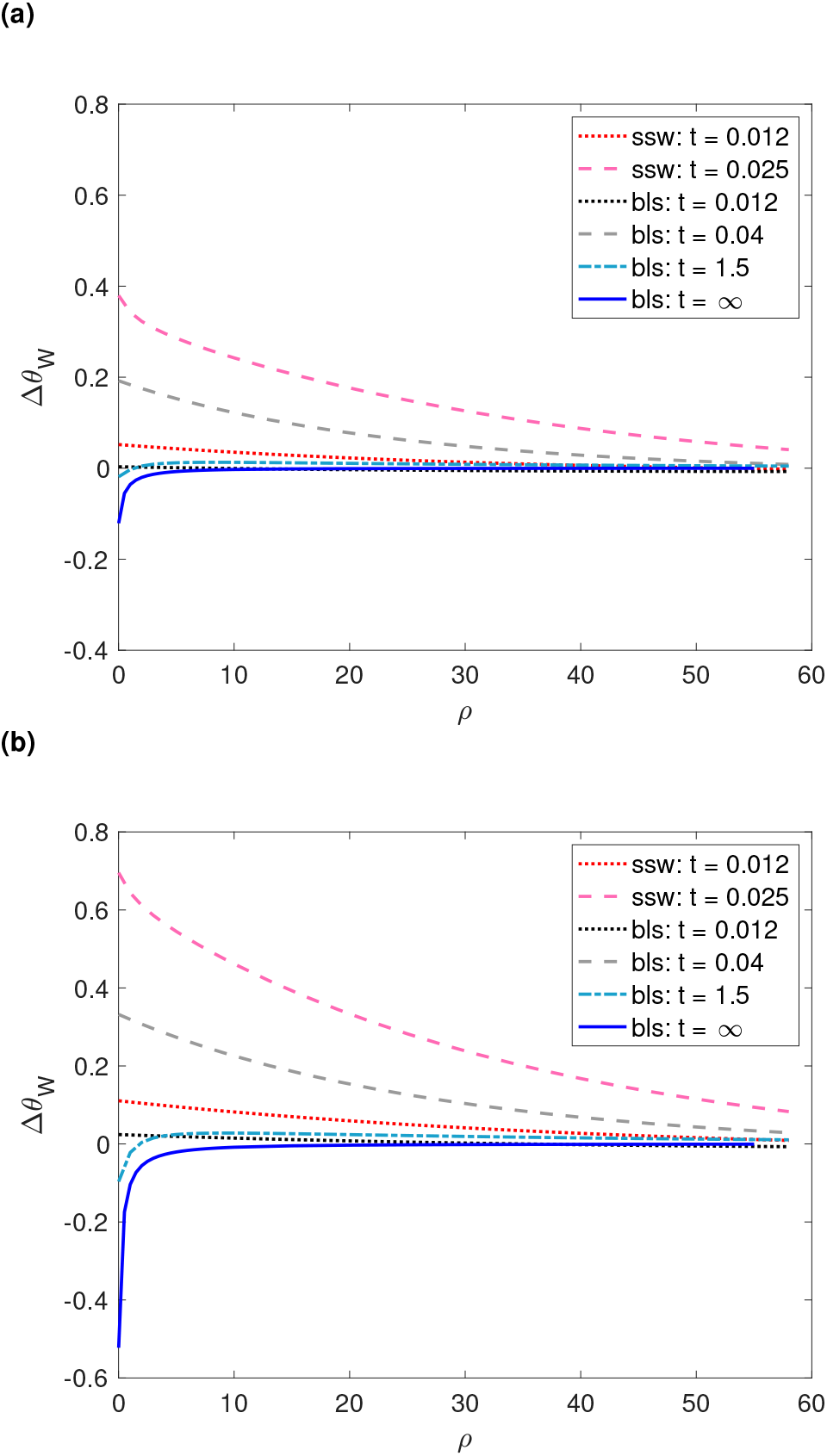
Δ*θ_W_* as a function of *ρ* and *t*. The two selection models are the same as those considered in Figure 9. “bls: *t* =∞” corresponds to the equilibrium under balancing selection. The sample size is 10 in (a) and 35 in (b).

It is informative to compare the three balancing selection models with *γ*_1_ = 500, but different equilibrium allele frequencies (Figure 6). The model with 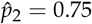 produces the strongest sweep-like signals (Figure 10 vs Figure S14). At the other extreme, the model with 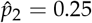 effectively emits no such signal (Figure S14). Thus, targets of recent balancing selection with larger 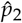 are easier to detect. However, for older targets of selection, the excess of intermediate frequency variant (i.e., negative Δ*θ_W_*) is most noticeable for selection targets with 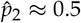 (Figure S14), making them the most amenable to detection. Altogether, it seems that balancing selection targets with low equilibrium allele frequencies (e.g., 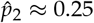) are difficult to identify regardless of their age.

### Simulations

We performed simulations with stochastic allele frequency trajectories at the selected site using `mbs`. The simulation method is similar to that described earlier (see also Supplement Text S.6). In Figure S15, *γ*_1_ = 500 and the equilibrium frequency of *A*_2_ is 0.75 (i.e., the same as Figure 9). The theoretical predictions for both the balancing selection and selective sweep models are highly accurate. In an additional experiment, we reduced *γ*_1_ to 20, but kept the equilibrium frequency of *A*_2_ at 0.75. This is to examine the robustness of our predictions against increased stochasticity induced by weaker selection. The results in Figure S16 suggest that our theory remains accurate for both models.

## Data availability

The methods for performing the calculations presented in this paper have been implemented in an R package named bls, and is available from http://zeng-lab.group.shef.ac.uk. In addition to the models considered here, the package can also obtain the total branch length and the SFS for (1) neutral models with changes in population size, (2) neutral models with two demes and changes in migration rates and/or deme sizes, and (3) isolation with migration models.

## Discussion

In this study, we have used the power and flexibility afforded by phase-type theory to study the effects of balancing selection on patterns of genetic variability and LD in nearby genomic regions. Our results go beyond previous attempts in that they provide a unifying framework for calculating important statistics for both equilibrium and nonequilibrium cases. In what follows, we discuss how our results can be used in data analyses and future method developments. We will also discuss the usefulness of phase-type theory in general.

### Accommodating other biological factors

Here we have only considered selection on an autosomal locus in a randomly mating population. However, our results can be readily extended to accommodate other important biological factors. Take self-fertilization as an example. Let *f* be the selfing rate and *F* = *f* / (2 – *f*) be the corresponding inbreeding coefficient. For this model, *N_e_* = *N*/ (1 + *F*), where *N* is the number of breeding individuals (Charlesworth 2009). Because selfing increases the frequency of homozygotes in the population, it reduces the effective frequency of recombination to *r_e_* = (1 – *F*)*r*, where *r* is the autosomal recombination rate in a random-mating population (Nordborg 1997; see Hartfield and Bataillon 2020 for a more accurate expression for *r_e_*). Finally, for the model of recent balancing selection, we also need to consider the effects of selfing on the frequency trajectory of *A*_2_. This can be achieved by replacing (22) with:

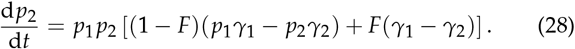

Other factors, including separate sexes, mode of inheritance (e.g., X-linkage vs autosomal), and background selection, can also be modelled (Charlesworth 2009; Vicoso and Charlesworth 2009; Glémin 2012; Charlesworth 2020a; Hartfield and Bataillon 2020).

### Detecting long-term balancing selection

We have examined two models of long-term balancing selection, one with a constant population size and the other with recent demographic changes. We confirm the well-known result that long-term balancing selection leads to elevated diversity, increased LD, and an excess of intermediate frequency variants in the SFS (Figures 2 - 4, 10; Charlesworth 2006; Fijarczyk and Babik 2015). Because the strength of these signals is weak except at sites very close to the locus under selection, they could be useful in pinpointing targets of balancing selection. On the other hand, we find that, under our two-allele model, these signals are strongest when the equilibrium frequencies of the selected variants are close to 50% (Figures 2 - 4, 10, and S14). This implies that genome scan methods are likely to be biased towards detecting selection targets where the selected variants are more common, which appears to be the case for some detection methods (Bitarello *et al*. 2018; Siewert and Voight 2020).

Our results can be used to improve existing methods for detecting balancing selection. For example, the *T*_1_ test by DeGiorgio *et al*. (2014), which has been shown to be among the most powerful, is based on *L*, the expected total branch length. The recursion equations DeGiorgio *et al*. (2014) used to obtain *L* assumes a constant population size. We can now relax this assumption by incorporating changes in population size. The increase in the strength of signals of long-term balancing selection after population size reduction (Figure 5b) points to the importance of incorporating non-equilibrium demographic dynamics, which may help to increase statistical power and reduce false positive rates. Nonetheless, the results presented in Figures 4 and 10 show that *L* does not capture all of the information about balancing selection. Instead, statistical power can be gained by making use of the SFS. This explains why the *T*_1_ test (based on *L*) is often less powerful than the *T*_2_ test (based on the SFS) (DeGiorgio *et al*. 2014). However, DeGiorgio *et al*. (2014) obtained the SFS via stochastic simulations, due to a lack of analytical methods. Here we have filled this gap. As above, it is of importance to extend the *T*_2_ test, so that it includes both the equilibrium and non-equilibrium models.

### Detecting recent balancing selection

It has long been suggested that signals generated by recent balancing selection should be similar to those generated by incomplete sweeps (Charlesworth 2006; Fijarczyk and Babik 2015). However, the allele frequency trajectories under these two models are similar only when the mutant allele is rather rare in the population (Figure 6). This period accounts for a small fraction of the time it takes to fix a positively selected mutation subject to a comparable level of selection. In addition, the rate of allele frequency change in this period is slower than when the mutant allele is more common. Combining these two factors, it is unsurprising that, at the time when the allele frequency trajectories under the two models start to diverge, neither model produces a noticeable effect on diversity patterns in nearby genomic regions (data not shown). Thus, this initial period of identity contributes very little signal.

After the initial period, the frequency of the positively selected mutation increases rapidly. In contrast, the rate of growth under the balancing selection model is much slower, especially when the equilibrium frequency of the mutant allele is low (Figure 6). Nonetheless, the increase in frequency of a recent balanced polymorphism does produce sweep-like diversity patterns. These include reductions in genetic variability, a skew towards high and low frequency derived variants in the SFS, and a build-up of LD between the selected and linked neutral sites (Figures 7 - 10). In addition, the maximum build-up of LD appears before the reduction in diversity levels and the distortion of the SFS peak, suggesting that these signals complement each other. Although these patterns are not as pronounced as those produced by sweeps of a comparable strength, we expect them to be detectable by methods designed for identifying sweeps (Booker *et al*. 2017; Pavlidis and Alachiotis 2017), as has been shown previously (Zeng *et al*. 2006). An open question is whether it is possible to distinguish between these two types of selection. On the other hand, because recent balancing selection causes diversity and LD patterns to be in a non-equilibrium state for a long period (Figures 10 and S14), it is unclear whether these patterns can be exploited for detecting selection targets.

Comparing the three balancing selection models with equilibrium allele frequencies 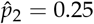, 0.5, and 0.75, respectively (Figure 6), mutations with 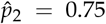 produce the strongest sweep-like patterns (e.g., Figure 9 vs Figure S13). They are probably the easiest to detect, although they may also be the most difficult to be distinguished from sweeps. On the other hand, although selection targets with 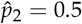 are not as easy to detect when they are young, they produce the strongest deviation from neutrality if they have been maintained for a sufficiently long period of time (Figures 2, 3, and S14), suggesting that they are most likely to be identified by methods for detecting long-term selection targets. Finally, it seems that selection targets with 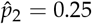 are the most difficult to detect regardless of the age of the mutant allele.

### Using phase-type theory to assess the accuracy of simpler approximations

We have shown the ease for which phase-type theory can be used to analyse complex models. In some cases, this can lead to simple analytic solutions (e.g., (7) and (8)). When explicit analytic solutions are difficult to obtain, phase-type theory can be useful in searching for simpler approximations. Take the model of recent balancing selection as an example. By using a large number of bins in the discretisation scheme (Figure S8), we can obtain results that are effectively exact. It is, however, impossible to write them as simple equations. Nonetheless, if we make an additional assumption that the recombination frequency between the selected locus and the neutral locus is not too high relative to the strength of selection, we can adopt the methods developed in Charlesworth (2020b) for selective sweeps, such that they can be used to obtain the expected pairwise coalescence time (see Supplementary Text S.8 for details).

We can assess the reliability of this approximation by comparing its results with those obtained using the phase-type method. As expected, the approximate results match the exact results closely when the recombination rate is low (e.g., *ρ* =1 in Figure 11). For higher recombination rates, the approximation under-estimates the diversity-reducing effect of the spread of *A*_2_. The main reason for this discrepancy is that the approximation assumes that the recombination rate is low, and the “sweep phase” is short. When these assumptions hold, once recombination during the sweep phase has moved a lineage from allelic class 2 to allelic class 1, back migration to allelic class 2 can be ignored. Although these assumptions work well for selective sweep models (Charlesworth 2020b), they are less suitable for the model of recent balancing selection, because the increase in allele frequency is much slower, leading to a longer sweep phase, and hence more opportunities for recombination. Thus, by preventing lineages from being moved back into allelic class 2, the approximation artificially slows down the rate of coalescence during the sweep phase, explaining the overestimation of pairwise coalescence time. Using results produced by phase-type theory as the baseline is desirable because, unlike stochastic simulations, these results are analytical, making comparisons straightforward and small differences easier to detect.

**Figure 11.**
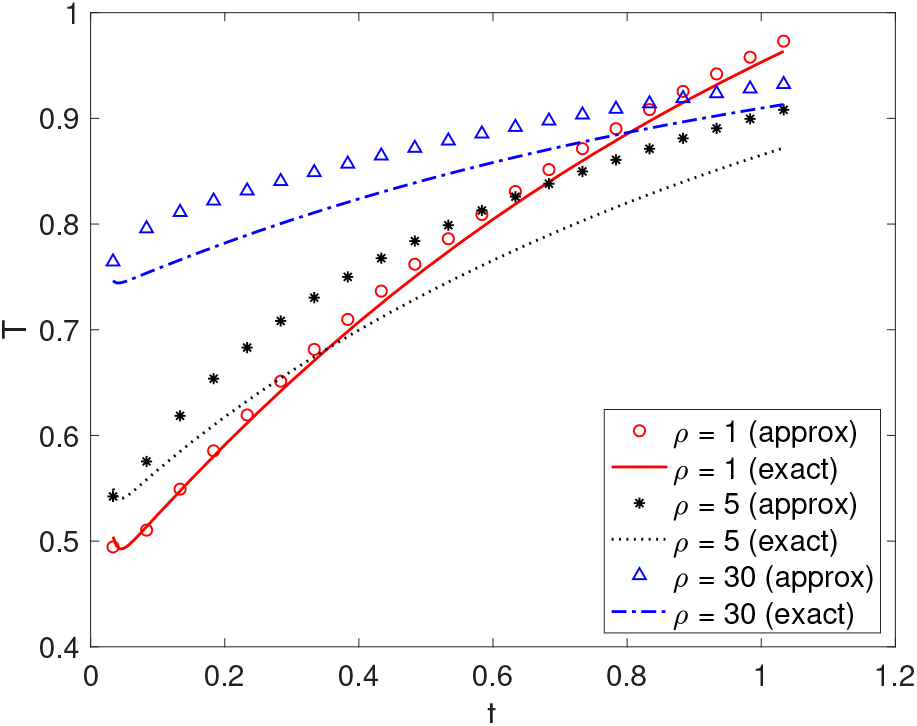
Comparing expected pairwise coalescence times obtained by phase-type theory (exact) and an approximation assuming low recombination rates. The model of recent balancing selection model has the following parameters: *γ*_1_ = 500 and 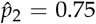 (i.e., the same as in Figures 7 - 10). *t* is the time since the arrival of *A*_2_. The discretisation scheme has *H* = 76 epochs. Details of the approximation are given in Supplementary Text S.8.

### Differences from previous studies and limitations

The equilibrium model of balancing selection has been analysed previously using coalescent theory (Hudson and Kaplan 1988; Nordborg 1997). Phase-type theory has allowed us to reproduce well known results (e.g., (8)). Additionally, it has made it feasible to obtain other important summary statistics (e.g., total branch length, LD and SFS) and introduce non-equilibrium scenarios (changes in population size or recent selection). Recently, Kern and Hey (2017) analysed a coalescent model with isolation and migration. Although the authors did not consider selection, the approach they used is related in that it involves performing calculations directly using the underlying continuous time Markov process. However, the results derived using our formulation is more compact (e.g., Theorem 1), which facilitates the accommodation of more complex situations (e.g., recent selection). Furthermore, we are able to obtain other useful results such as the second moment of the mean time to MRCA (Theorem 2 in Supplementary Text S.7).

A limitation of the phase-type approach is that the size of the state space increases quickly with the sample size, meaning that the computational cost will become too high for large samples. However, there is evidence that samples with as few as 20 alleles, which is computationally feasible using our approach, offer sufficient statistical power for detecting balancing selection (Siewert and Voight 2017; Bitarello *et al*. 2018). More importantly, our method provides a way of analysing complex models, which will help us to understand their properties. This may in turn enable us to obtain computationally more efficient approximations, as shown in the previous section. Finally, although the speed of forward simulators has improved significantly (Haller and Messer 2019), the phase-type approach is still much faster for moderate sample sizes. This is because, for a given set of parameters, we only need to perform the calculation once to obtain, for instance, the expected total branch length. In contrast, obtaining this quantity accurately using simulations requires at least tens of thousands of replicates. Simulations are, however, highly flexible and can be used to study models that are too difficult to analyse mathematically. Thus, both mathematical modelling and simulations are important.

### Applying phase-type theory to other population genetic mod-els

Phase-type theory can be applied to many different models in population genetics. For example, Hobolth *et al*. (2019) used a time-homogeneous version of the theory to study the standard Kingman’s coalescent with and without recombination, coalescent models with multiple mergers, and coalescent models with seed banks. They showed the ease for which useful results can be obtained (e.g., all the moments of the pairwise coalescence time, the covariance in coalescence times between two linked loci, or the SFS). By extending the framework to non-equilibrium cases (see Theorem 1, Corollary 1 in Supplementary Text S.5, and Theorem 2 in Supplementary Text S.7), we make this approach applicable to a yet larger class of models. For instance, we can introduce population size fluctuations into the models considered by Hobolth *et al*. (2019). Even for models that have been analysed before using other approaches (e.g., Matuszewski *et al*. 2017), it is worth exploring whether the new theory provides a better alternative, both in terms of ease of analysis and numerical stability of the resulting method, which may be beneficial for parameter estimation purposes (e.g., Kern and Hey 2017).

The phase-type approach may be particularly useful for models that involve selection on a single locus at which the frequencies of the selected variants change deterministically (Maynard Smith and Haigh 1974; Kaplan *et al*. 1988; Coop and Ralph 2012). These include the balancing selection models considered here, selective sweep models (Barton 1998; Kim and Stephan 2002; Kim and Nielsen 2004; Ewing *et al*. 2010; Charlesworth 2020a; Hartfield and Bataillon 2020), soft sweeps caused by recurrent mutation or migration (Pennings and Hermisson 2006), incomplete sweeps (Vy and Kim 2015), and recurrent sweeps (Kaplan *et al*. 1989; Kim 2006; Campos and Charlesworth 2019).

Here, we have briefly considered selective sweep models with semi-dominance and compared it to the corresponding balancing selection model (see (24) and Figures 6, 8 – 10). In a related study, we will use the phase-type approach to investigate some of the sweep models listed above more systematically (K. Zeng and B. Charlesworth, *in preparation*). Because we can use phase-type theory to obtain exact solutions, it provides a convenient way to determine the accuracy of existing approximations. For instance, for the sweep model with semi-dominance, a widely-used approximation assumes that there is no coalescence during the sweep phase, such that the gene tree for a set of alleles sampled immediately after a sweep has a simple “star shape” (Maynard Smith and Haigh 1974; Barton 2000; Durrett and Schweinsberg 2004). However, a recent study of the pair-wise coalescence time suggests that this approximation can be rather inaccurate when the ratio of the recombination rate to the selection coefficient is high (Charlesworth 2020b). It is important to also assess the effect of this simplifying assumption on the SFS, given that both nucleotide site diversity and the SFS are informative when it comes to estimating the strength and prevalence of (recurrent) sweeps (Corbett-Detig *et al*. 2015; Elyashiv *et al*. 2016; Booker *et al*. 2017; Comeron 2017). In addition, we can also explore the joint effects of recurrent sweeps and recent population size changes. These are not well understood, but are important for estimating the relative importance of background selection and recurrent sweeps in shaping genome-wide patterns of variability (e.g., Johri *et al*. 2020).

## Supplementary text

### S.1 Definition of the backward mutation rate

Consider the equilibrium model of balancing selection. The equilibrium frequency of the selected variant *A_i_* is 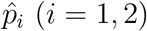. During reproduction, an *A_i_* allele mutates to *A_j_* with probability *u_ij_* per generation. Forward in time, after reproduction, the frequency of *A_i_* becomes 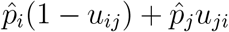, where the second term is the frequency of mutants amongst all *A_i_* alleles. Going backward in time, the backward mutation rate is the probability that an *A_i_* allele in the post-reproduction population is descendant from an *A_j_* parent via mutation:

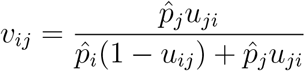

Assuming that *u_ij_* is much smaller than both 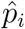 and 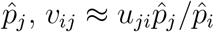, which is the same as (6) in Kaplan *et al*. (1988).

### S.2 Calculating the Green’s matrix

Let {*X_t_*}_*t*≥0_ be a Markov jump process with sub-intensity matrix ***S***. That is, *X_t_* = *i* means that the process is in state *i* at time *t*. Let 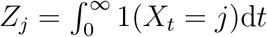 denote the time spent in state *j* before absorption, where 1(*X_t_* = *j*) = 1 if *X_t_* = *j*, and 0 otherwise. The expected time spent in state *j* given initial state *i* is then

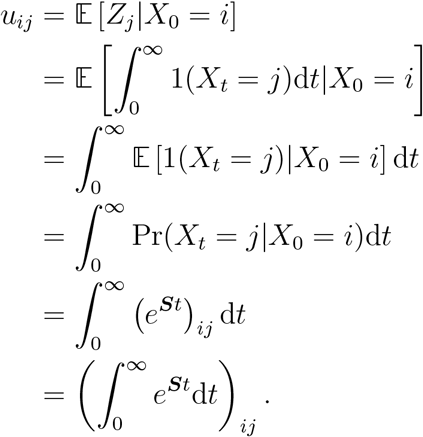

To complete the proof, we note that

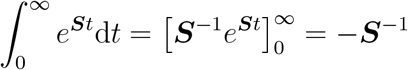

where we have used the property that (*e^**St**^*)_*ij*_ → 0 for *t* → ∞.

### S.3 The intensity matrix for calculating the total branch length of a sample size of three

***S***_2_ and ***s***_2_ in (12) are the same as the corresponding elements defined in (3).

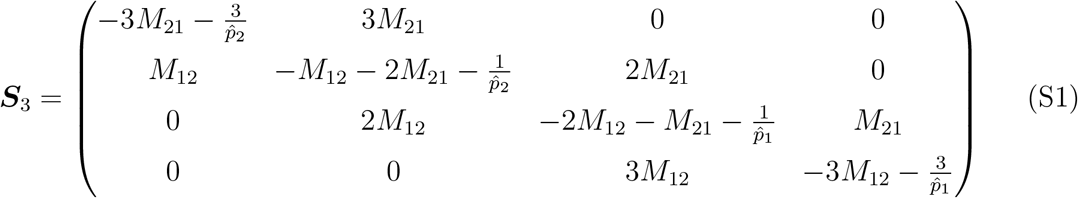

and

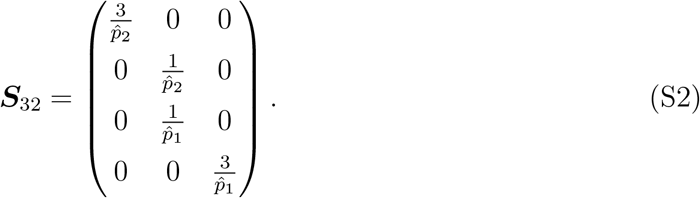

### S.4 The intensity matrix for calculating the SFS for a sample size of three

The sub-matrices in (12) for the model leading to Table 1 are given below.

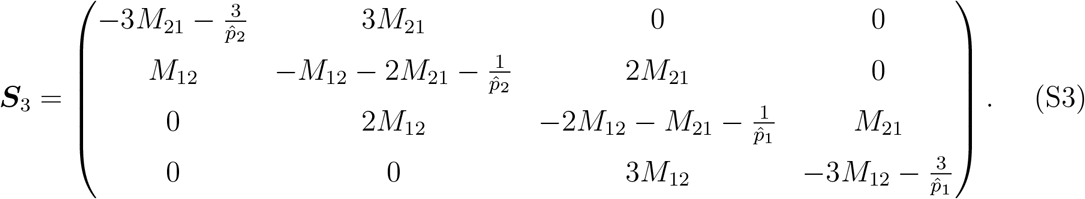

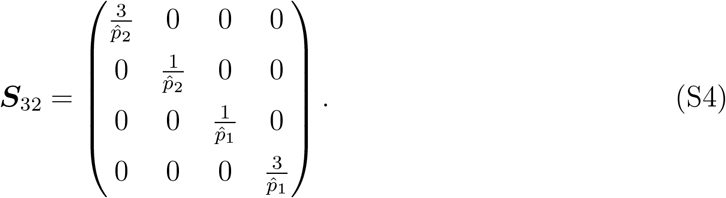

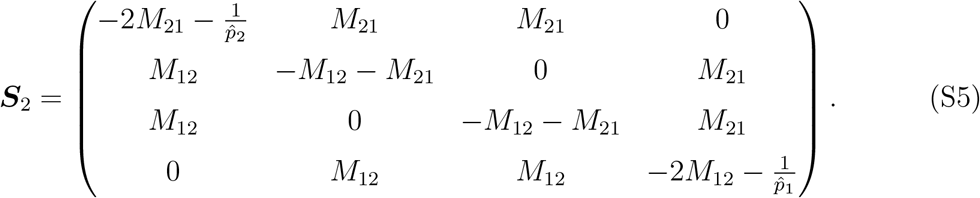

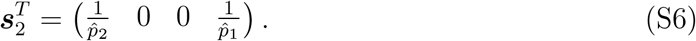

### S.5 A non-equilibrium phase-type model

Consider a continuous time Markov chain with finite state space {1, 2,…, *K, K* + 1}, where states 1,…, *K* are transient, and state *K* + 1 is absorbing. It is assumed that the time interval [0, ∞) is subdivided into *H* non-overlapping epochs. The duration of epoch *h* is [*t*_*h*–1_, *t_h_*), where 1 ≤ *h* ≤ *H*, *t*_0_ = 0, and *t_H_* = ∞. The intensity matrix for epoch *h* is constant and takes the form:

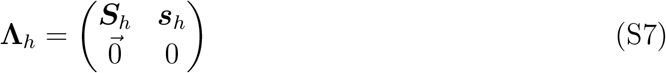

where ***S**_h_* the *K*-by-*K* sub-intensity matrix, and ***s**_h_* is the *K*-by-1 exit rate vector.

Define

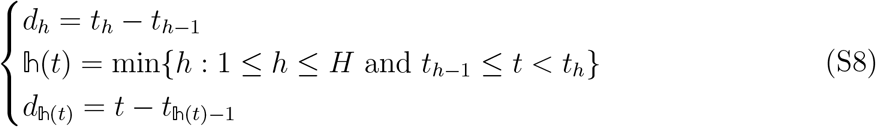

The transition probability between time 0 and time *t* is given by:

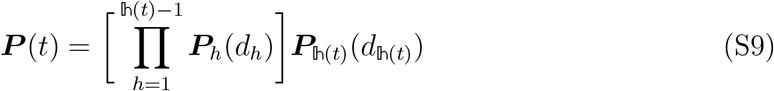

where ***P**_h_*(*t*) is the transition matrix for epoch *h*. Note that the matrices do not commute. So the multiplication should be carried out from left to right, according to the chronological order of the epochs. From standard Markov chain theory, we know that:

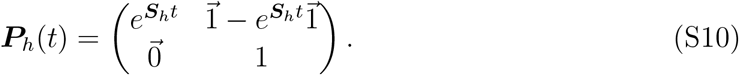

Define

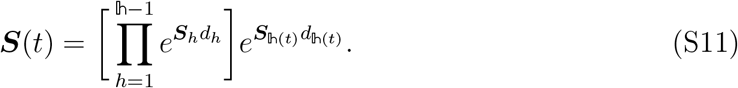

We can rewrite (S9) in a more compact form:

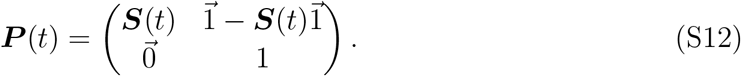

The probability that the process jumps to the absorbing state in the time interval [*t, t*+*dt*) is given by:

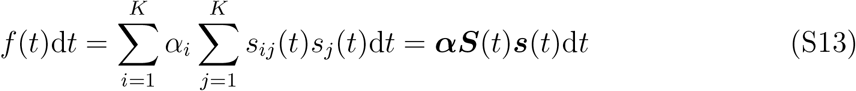

where ***α*** is the initial probability vector, *s_ij_*(*t*) are elements of ***S***(*t*), and *s_j_*(*t*) are elements of 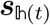, the exit rate vector at time *t*. The Laplace transform of *f*(*t*) is defined as:

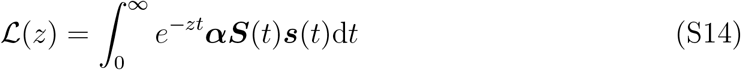

for *z* ≥ 0. Noting that 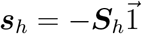 and substituting (S11) into (S14) leads to:

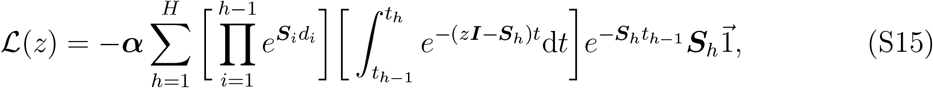

where ***I*** is the identity matrix. To evaluate the integral, we define ***A**_h_*(*z*) = ***A**_h_* = – (*z**I*** – ***S**_h_*). Because all eigenvalues of ***A**_h_* have strictly negative real parts (Hobolth *et al*., 2019), 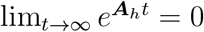. We obtain:

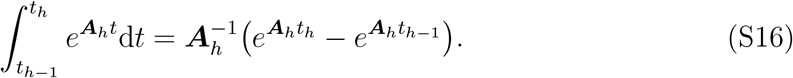

Taking the derivative with respect to *z*, we obtain:

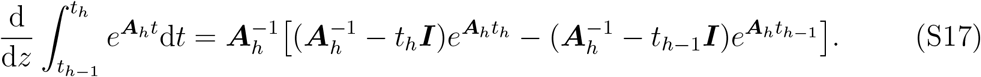

Noting that the mean time to absorption is given by 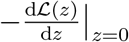 and that ***A**_h_*(0) = ***S**_h_*, we have:

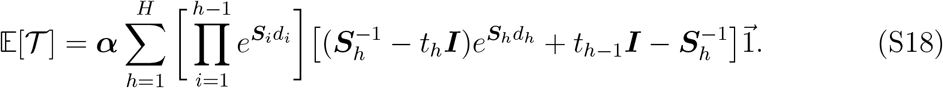

Expanding the equation and removing terms that cancel each other, we arrive at Theorem 1. To facilitate further discussion, we state this Theorem in a slightly different way:

#### Corollary 1.

*Let **α*** = (*α*_1_,…, *α_K_*), *where α_i_ is the probability that the initial state is i and* 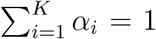. *Let* 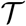 *be a random variable representing the time to absorption. We have:*

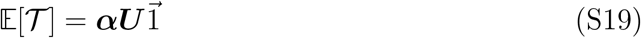

*where*

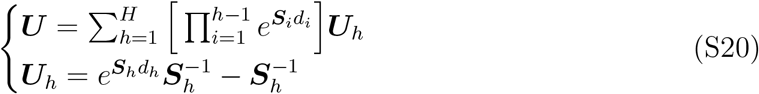

*and e^**S**_h_d_h_^* = 0 *if d_h_* = ∞.

We have also derived an expression for the second moment of 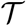 in Theorem 2 in Supplementary Text S.7.

Let *u_ij,h_* represent the elements of ***U**_h_. U_ij,h_* is the amount of time the process spends in state *j* during [*t*_*h*–1_, *t_h_*) given that it is in state *i* at time *t*_*h*–1_. That is, ***U**_h_* is the Green’s matrix for the *h*-th epoch. Also note that element *i* in the vector 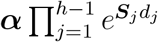 gives the probability that the process is in state *i* at time *t*_*h*–1_. Thus, Corollary 1 shows that, under this stepwise model, the Green’s matrix for the entire process ***U*** is the weighted average of the Green’s matrices of all the constituent epochs.

As noted in the main text, the expectation of both *L*_*n*_1_, *n*_2__ and *ϕ*^(*n*_1_, *n*_2_)^ can be written in the form ***αUD***. Let *Y* represent either of these two random variables. Corollary 1 tells us that:

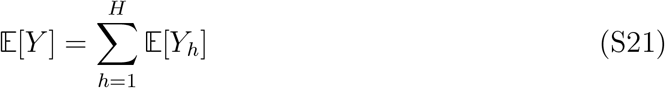

where

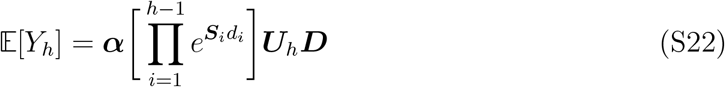

which is the expected contribution from epoch *h*.

We have so far assumed that the state space is the same across epochs. This restriction can be relaxed. Let the size of the state space in epoch *h* be *K_h_*. Let ***E***_*h*–1, *h*_ be a *K*_*h*–1_-by-*K_h_* matrix that defines the mapping of the states from epoch *h* – 1 to epoch *h* (*h* = 1,…, *H* and *E*_01_ = ***I***, the identity matrix). Corollary 1 holds if we replace 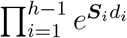 by 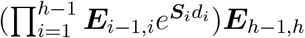. For (S21), we additionally need to replace ***D*** by an epoch-specific ***D**_h_*.

### S.6 Coalescent simulation with stochastic allele frequency trajectories

The theory developed in the main text assumes that the allele frequencies of the variants at the selected locus change deterministically over time. In reality, frequencies of the selected variants fluctuate because of random genetic drift. To investigate the effects of stochastic allele frequency fluctuation on the accuracy of our model predictions, we conducted simulations with stochastic allele frequency trajectories. Each simulation replicate contained two steps: (1) forward simulation to obtain allele frequency trajectories for the selected variants given the demographic history; (2) coalescent simulation for a sample of *n* alleles at a linked neutral site, conditioning on the trajectories obtained in step 1. The forward simulation step was based on a custom R script, which is available on request. The coalescent simulation step was performed using `mbs` (Teshima and Innan, 2009).

#### Forward simulations

Consider a Wright-Fisher population. The variants at the focal selected site are represented by *A*_1_ and *A*_2_, respectively. The fitnesses of *A*_1_*A*_1_, *A*_1_*A*_2_, and *A*_2_*A*_2_ are denoted by *w*_11_, *w*_12_, and *w*_22_, respectively. The model has the following life cycle: mutation, random mating, selection, and sampling. The mutation rate from *A*_1_ to *A*_2_ is *u*_12_ per generation, and that in the opposite direction is *u*_21_. Note that reversible mutation between *A*_1_ and *A***2** is included in the life cycle to make the model consistent with `mbs`.

Let the frequency of *A*_2_ in generation *t* be *p*_2_. After mutation, it becomes 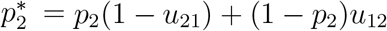. After random mating and selection, its frequency becomes

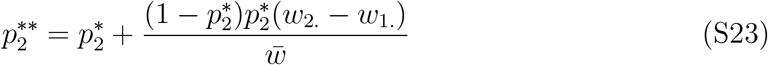

where 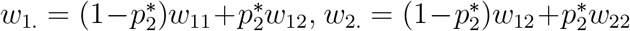, and 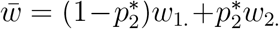. Let the population size of generation *t* + 1 be *N*_*t*+1_. We draw a random number *X* from the binomial distribution with parameters 2*N*_*t*+1_ and 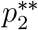. The frequency of *A*_2_ in generation *t* + 1 is therefore *X*/(2*N*_t+1_). Thus, this approach can accommodate arbitrary changes in population size.

#### Models with strong balancing selection and changes in population size

This set of simulations are intended to check whether the results presented in Figures 5a and 5b are robust to stochastic fluctuation in the frequencies of the selected variants. We assume overdominance with *w*_11_ = 1 – *s*_1_, *w*_12_ = 1, and *w*_22_ = 1 – *s*_2_. The sample size was *n* = 20. Because *L* (the total branch length) is insensitive to the equilibrium frequencies of *A*_1_ and *A*_2_ when the sample size is not small (see Figure 4), we assumed a symmetric selection model with *s*_1_ = *s*_2_ = *s* (i.e., the equilibrium frequencies of *A*_1_ and *A*_2_ are 50%). To simulate the population expansion model presented in 5a, we assumed that *N*_*e*,1_ = 20, 000 (the effective population size of the current epoch) and *N*_*e*,2_ = 2, 000 (the effective population size of the ancestral epoch). For the population reduction model in 5b, we used *N*_*e*,1_ = 2, 000 and *N*_*e*,2_ = 20, 000.

For both demographic models, we assumed that the frequency of *A*_2_ at the beginning of the simulations was 50%. We then allowed the population to evolve forward in time for 50*N*_*e*,2_ generations. The population size was then changed to *N*_*e*,1_. We let the population evolve for another 2*N*_*e*,1_*t* generations, where *t* is the time parameter shown in Figure 5.

If either *A*_1_ or *A*_2_ was lost before the end of the simulation, the process was restarted. The allele frequency trajectory obtained was then passed onto `mbs` to obtain simulated sequence polymorphism data at a linked neutral site. *L* was estimated using *S*/*θ*, where *S* is the observed number of segregating sites, θ = 2*N*_*e*,1_*u*, and *u* is the neutral mutation rate.

#### Recent balancing selection and selective sweeps

Here we set out to test whether phase-type theory can produce accurate predictions about the SFS. As in Figure 10 in the main text, we considered overdominance with fitnesses *w*_11_ = 1 − *s*_1_, *w*_12_ = 1, and *w*_22_ = 1 − *s*_2_. The corresponding sweep model has fitnesses *w*_11_ = 1, *w*_12_ = 1 + *s*_1_, and *w*_22_ = 1 + 2*s*_1_. In the forward simulations, we assumed that *A*_2_ was the mutant allele, and it appeared as a single copy in the population when it first arose. We then allowed its frequency to evolve. We set *t* = 0 when the frequency of *A*_2_ exceeded ε = 1/*γ*_1_ for the first time, where γ_1_ = 2*Ns*_1_ and *N* is the effective population size. We allowed the forward simulation to continue until, time *t* (in units of 2*N* generations). The trajectory of *A*_2_ was then used by `mbs` for obtaining samples at a linked neutral site. The unfolded SFS as defined in the main text at the neutral site was estimated using data from 10,000 replicates.

### S.7 The second moment of the mean time to absorption

The second moment of 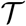 is given by 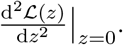 The second derivative with respect to *z* for the integral in (S16) reads:

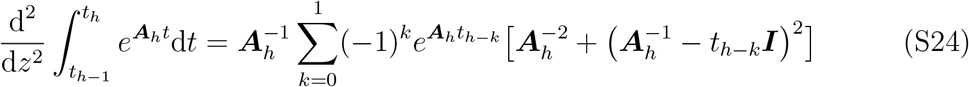

Substituting (S24) into (S15) leads to the following result.

#### Theorem 2.

*The second moment of the mean time to absorption*, 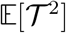, *is given by:*

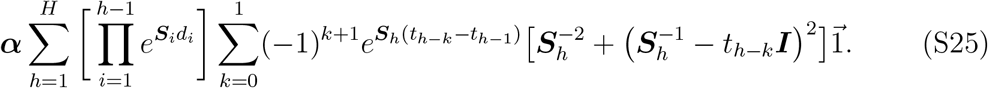

### S.8 Approximating the expected pairwise coalescence time under the model of recent balancing selection

As in the main text, we assume that a new allele *A*_2_ has arisen by mutation, and has spread to a frequency 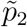 that is close to its equilibrium value under balancing selection, which is 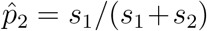 with heterozygote advantage. Providing that the recombination rate is not too high relative to the strength of selection, the expected coalescence time for a pair of *A*_2_ alleles sampled at frequency 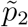 can be obtained from Equations 9, 10, 11a and A1-A3 of Charlesworth (2020b), where Δ*π* in his Equation 11a is equivalent to the reduction in the mean pairwise coalescence time relative to the neutral value of 2*N_e_* generations. To obtain △*π*, 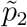 replaces *q*_2_ in Equations 9, 10 and A1-A3 of Charlesworth (2020b), where the selection parameters in Equations A1-A3 are *γ* = 2*N*_e_*s*_1_, *a* =1, and *b* = -(*s*_1_ + *s*_2_)/*s*_1_. At the time when 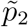 is reached, the values of the expected coalescent times (on the timescale of 2*N_e_* generations) for a pair of *A*_1_ alleles is approximately equal to 1.

In addition, the possibility that a recombination event introduces the neutral site from an *A*_1_ allele onto an *A*_2_ background, thereby reducing the initial divergence at the neutral site between an *A*_1_ and *A*_2_ pair, is modelled by using Equation A3a of Charlesworth (2020b) with *q*_2_ replaced with 1-*p*_2_ and *q* with 1 – *ϵ*, to yield a probability of an *A*_1_ to *A*_2_ recombination event of *P*_*r*1_. In addition, the selection parameters *a* and *b* should be replaced with *a* + *b*, and −*b*, respectively. It is assumed that such a recombination event is followed by coalescence with a non-recombined neutral site associated with *A*_2_, with a coalescence time equal to the duration of sweep, *t_s_*, as given by 23 with 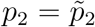. The divergence between an *A*_1_ and *A*_2_ pair at the time of sampling is then given by 1-*P*_*r*1_(1–*t_s_*).

A simple way to obtain the pairwise coalescence times at an arbitrary time after the allele frequency 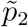 has been reached is to consider the recursion relations for the corresponding pairwise expected diversity measures with a neutral mutation rate of *u* under the infinite sites mutation model and assuming that the frequency of *A*_2_ remains close to its equilibrium value. The scaled mutation rate in the absence of selection, *θ* = 2*N_e_u*, is sufficiently small that second-order terms in *θ* can be neglected (Malécot, 1969, p. 40; Wiehe and Stephan, 1993, Equation 6a). Writing *π_ij_* for the expected diversity for a pair of alleles *A_i_* and *A_j_*, and using primes for their values in a new generation, and neglecting second-order terms, we have:

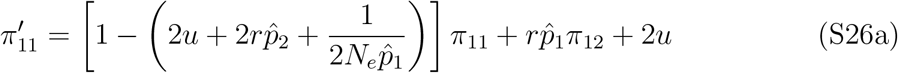

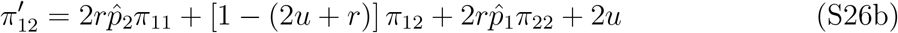

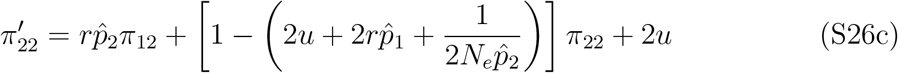

The coefficients of the *π_ij_* in these equations provide the corresponding coefficients for the recursions of the deviations of the *π_ij_* from their equilibrium values, thereby eliminating the term in 2u on the right-hand sides of the equations. If the *π_ij_* are scaled relative to their expected value 2*θ* in the absence of selection, and *u* is set arbitrarily close to zero, solving for equilibrium gives *π_ij_* values relative to 2*θ* that are equivalent to the equilibrium coalescent times given by (8), as can be verified by direct calculation.

By setting *u* to zero in (S26), and using the scaled the *π_ij_*, we thus obtain a recursion for the deviations from equilibrium of the corresponding expected pairwise coalescence times on the timescale of 2*N_e_* generations. While it is possible in principle to diagonalize the relevant matrix, and express the solution for an arbitrary time after reaching 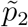 in term of its eigenvalues and eigenvectors, in practice it is simpler to iterate the matrix with assigned numerical values of the parameters. In order to save computation time, a relatively small value of *N_e_* can be used, and the recombination parameters rescaled accordingly to represent a much larger *N_e_* with the same value of *ρ* = 2*N_e_r*. The initial relative values of *π*_11_, *π*_12_, and *π*_22_ are 1, 1, and 1 – Δ*π*.

## 1 Supplementary table

**Table S1:** Robustness of the theoretical predictions with respect to selection intensity

| Expansion with $N_{e,1}/N_{e,2} = 10$ | | | | Reduction with $N_{e,1}/N_{e,2} = 0.1$ | | | |
| --- | --- | --- | --- | --- | --- | --- | --- |
| $\rho$ | Theory | $\gamma_{min} = 250$<br>Mean (SE) | $\gamma_{min} = 20$<br>Mean (SE) | $\rho$ | Theory | $\gamma_{min} = 250$<br>Mean (SE) | $\gamma_{min} = 20$<br>Mean (SE) |
| 0.05 | 38.40 | 38.58 (0.34) | 37.35 (0.33) | 0.05 | 101.83 | 101.16 (0.66) | 100.03 (0.65) |
| 0.10 | 21.08 | 21.03 (0.19) | 20.52 (0.19) | 0.10 | 81.53 | 80.70 (0.50) | 80.57 (0.49) |
| 0.20 | 11.79 | 11.62 (0.10) | 11.45 (0.10) | 0.15 | 74.51 | 72.51 (0.45) | 72.64 (0.46) |
| 0.40 | 6.97 | 6.89 (0.06) | 6.87 (0.06) | 0.20 | 70.89 | 69.62 (0.43) | 69.40 (0.43) |
| 0.80 | 4.52 | 4.49 (0.04) | 4.42 (0.04) | 0.25 | 68.67 | 67.95 (0.42) | 67.51 (0.42) |

The models used in the simulations are the same as those described in Figures S6 and S7. In both cases, samples were taken from the population at time $t = 0.1$ after the change in population size, where $t$ is expressed in units of $2N_{e,1}$ generations. We included neutral sites with different scaled recombination frequencies ( $\rho$ ) with the selected site. The mean and standard error of the total branch length $L$ for a sample of 20 alleles were estimated using 10,000 rounds of simulations.

## 2 Supplementary figures

**Figure S1:**
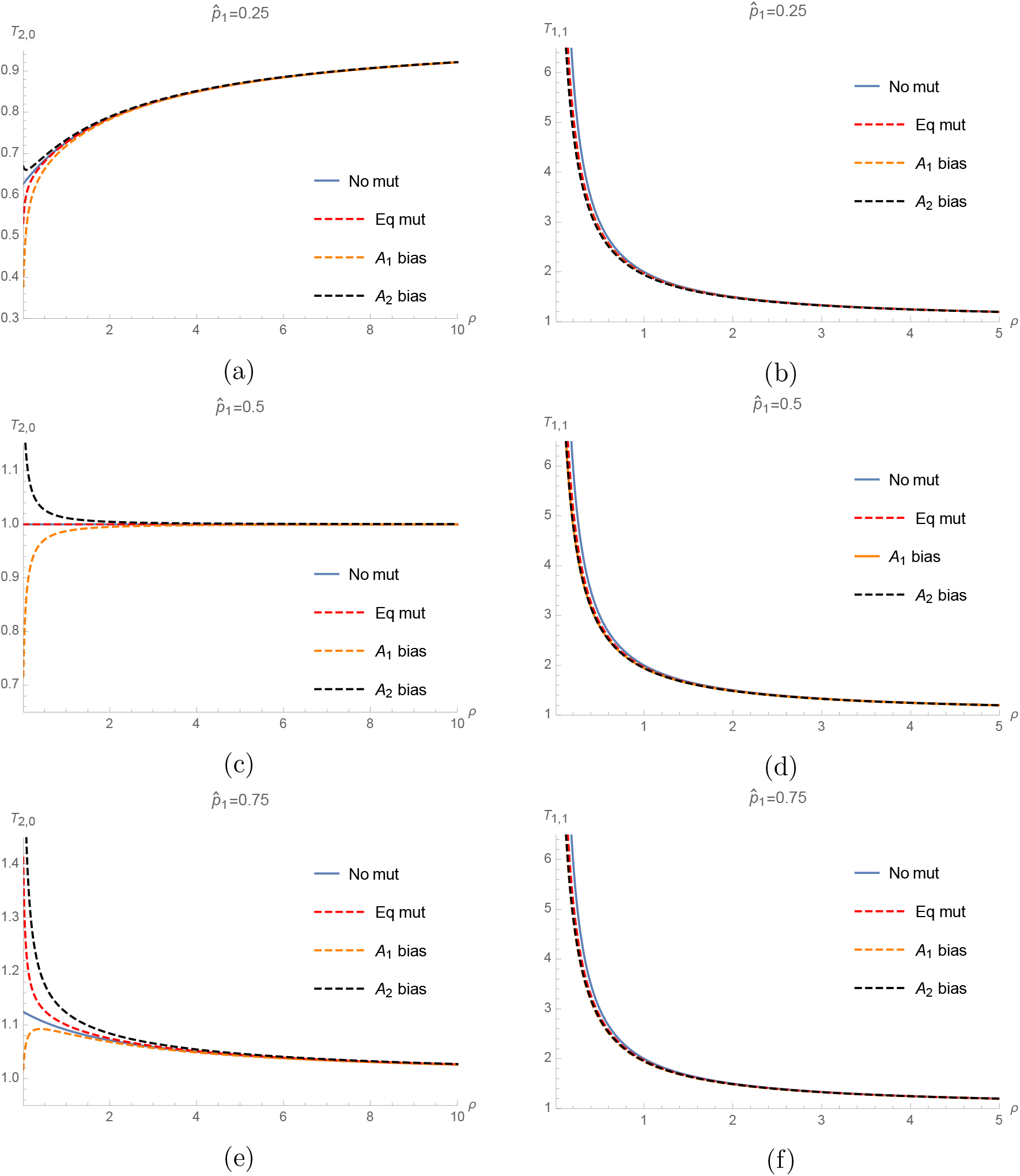
Expected coalescence time for a pair of alleles as a function of *ρ*. The selected alleles *A*_1_ and *A*_2_ are at equilibrium frequencies 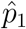 and 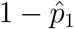. “No mut”means *μ_ij_* = 0 (i.e., (8)). “Eq mut” means *μ_ij_* = 0.02. “*A*_1_ bias” means *μ*_12_ = 0.01 and *μ*_21_ = 0.05. “*A*_2_ bias” means *μ*_12_ = 0.05 and *μ*_21_ = 0.01. The scales of the axes are different.

**Figure S2:**
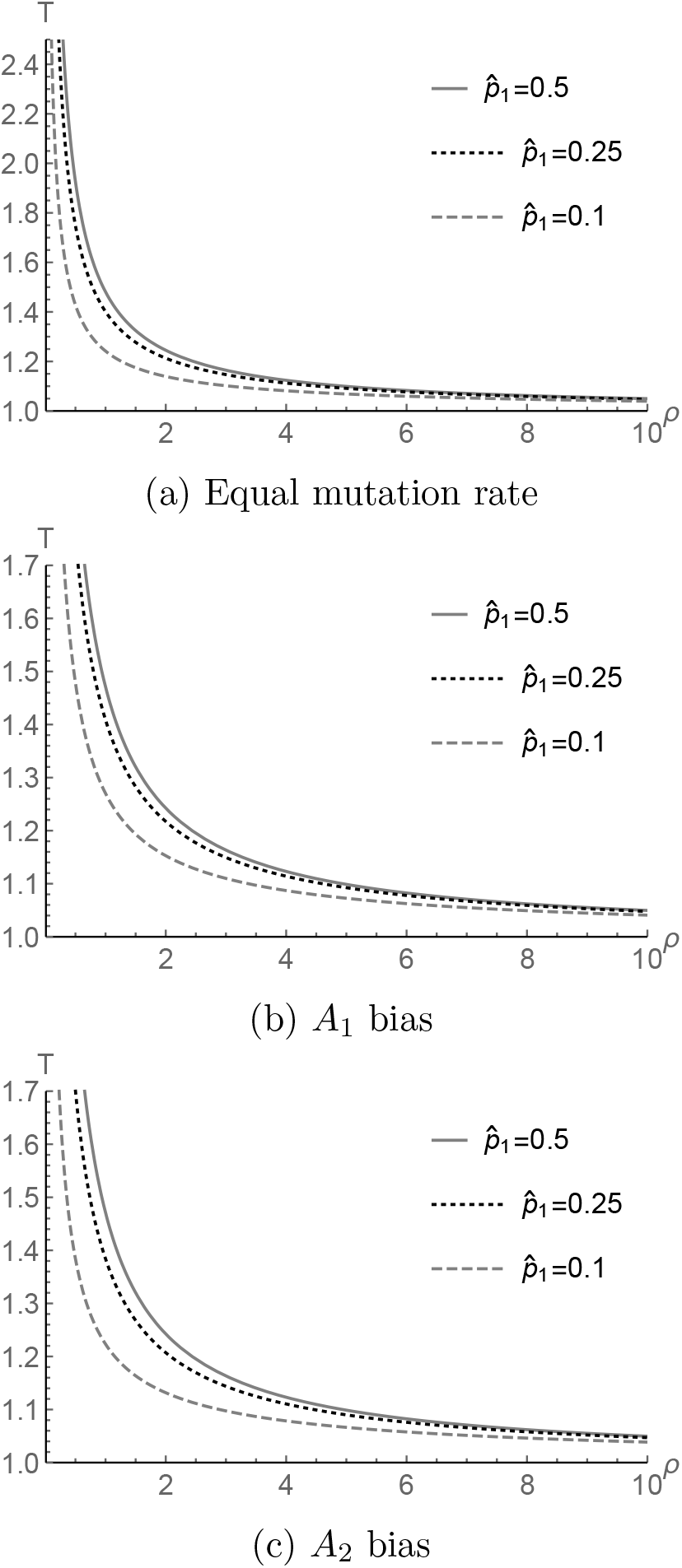
Expected coalescence time for a pair of alleles as a function of *ρ*. The selected alleles *A*_1_ and *A*_2_ are at equilibrium frequencies 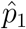 and 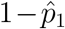. “Equal mutation rate” means *μ_ij_* = 0.02. “*A*_1_ bias” means *μ*_12_ = 0.01 and *μ*_21_ = 0.05. “*A*_2_ bias” means *μ*_12_ = 0.05 and *μ*_21_ = 0.01.

**Figure S3:**
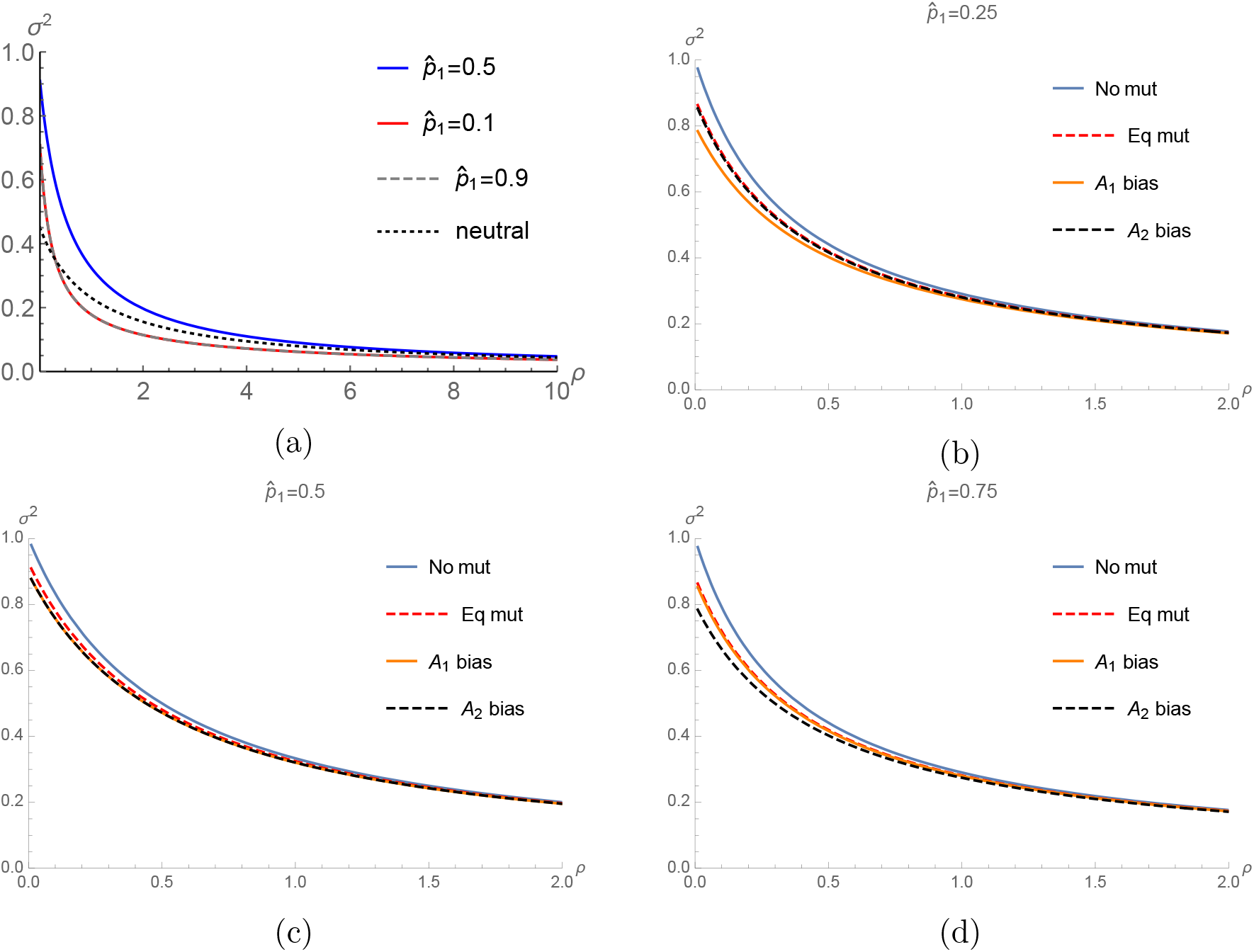
The level of LD between the selected and neutral loci as a function of *ρ*. In (a), the mutation rates between *A*_1_ and *A*_2_ are *μ*_12_ = *μ*_21_ = 0.02. In (b) - (d), for a given 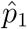, different mutation rates are considered. “No mut” means *μ_ij_* = 0. “Eq mut” means *μ_ij_* = 0.02. “*A*_2_ bias” means *μ*_12_ = 0.01 and *μ*_21_ = 0.05. “*A*_2_ bias” means *μ*_12_ = 0.05 and *μ*_21_ = 0.01.

**Figure S4:**
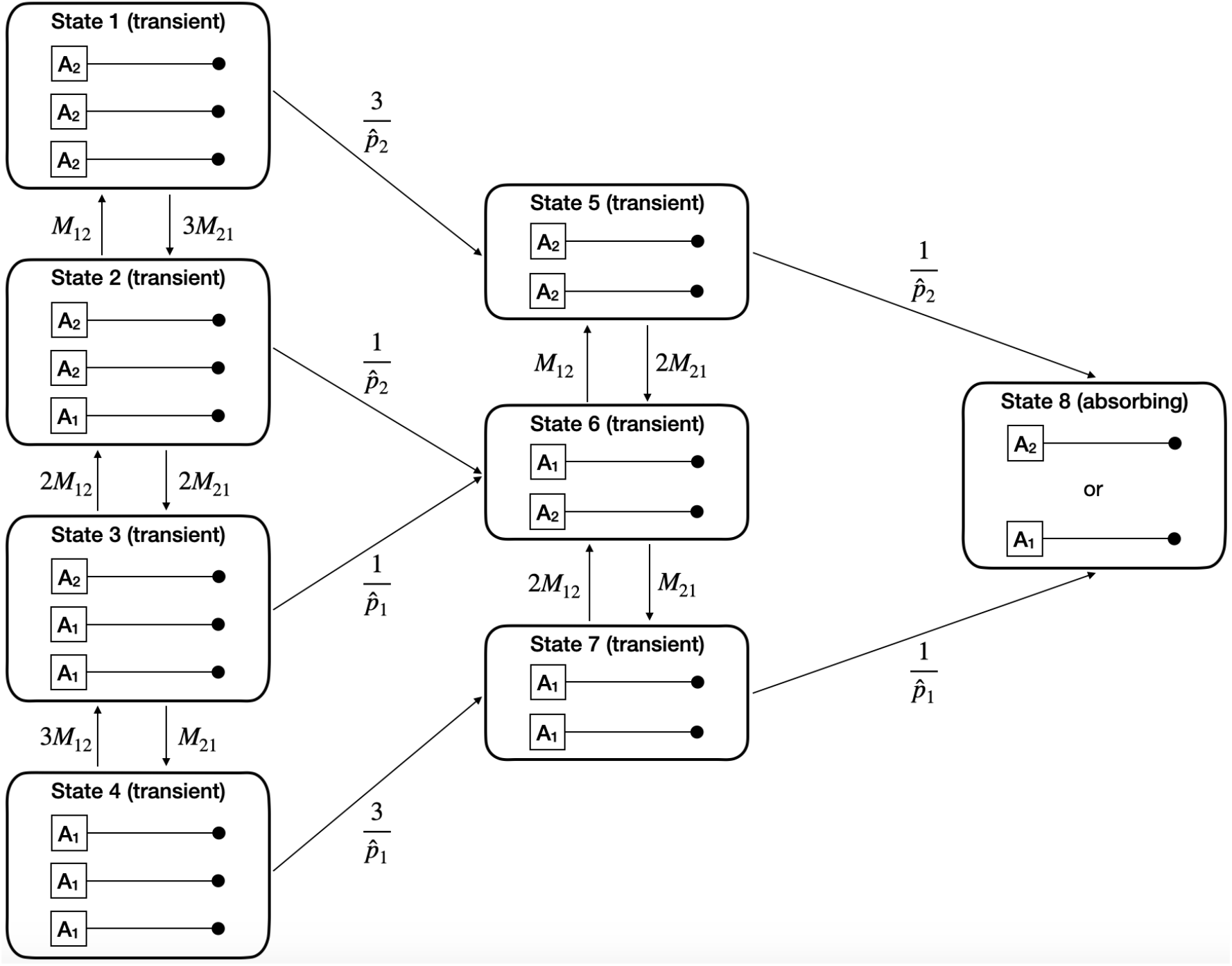
Transition rates between the states of the equilibrium balancing selection model for a sample of size three. Time is scaled in units of 2*N_e_* generations. The neutral locus is represented by a black dot.

**Figure S5:**
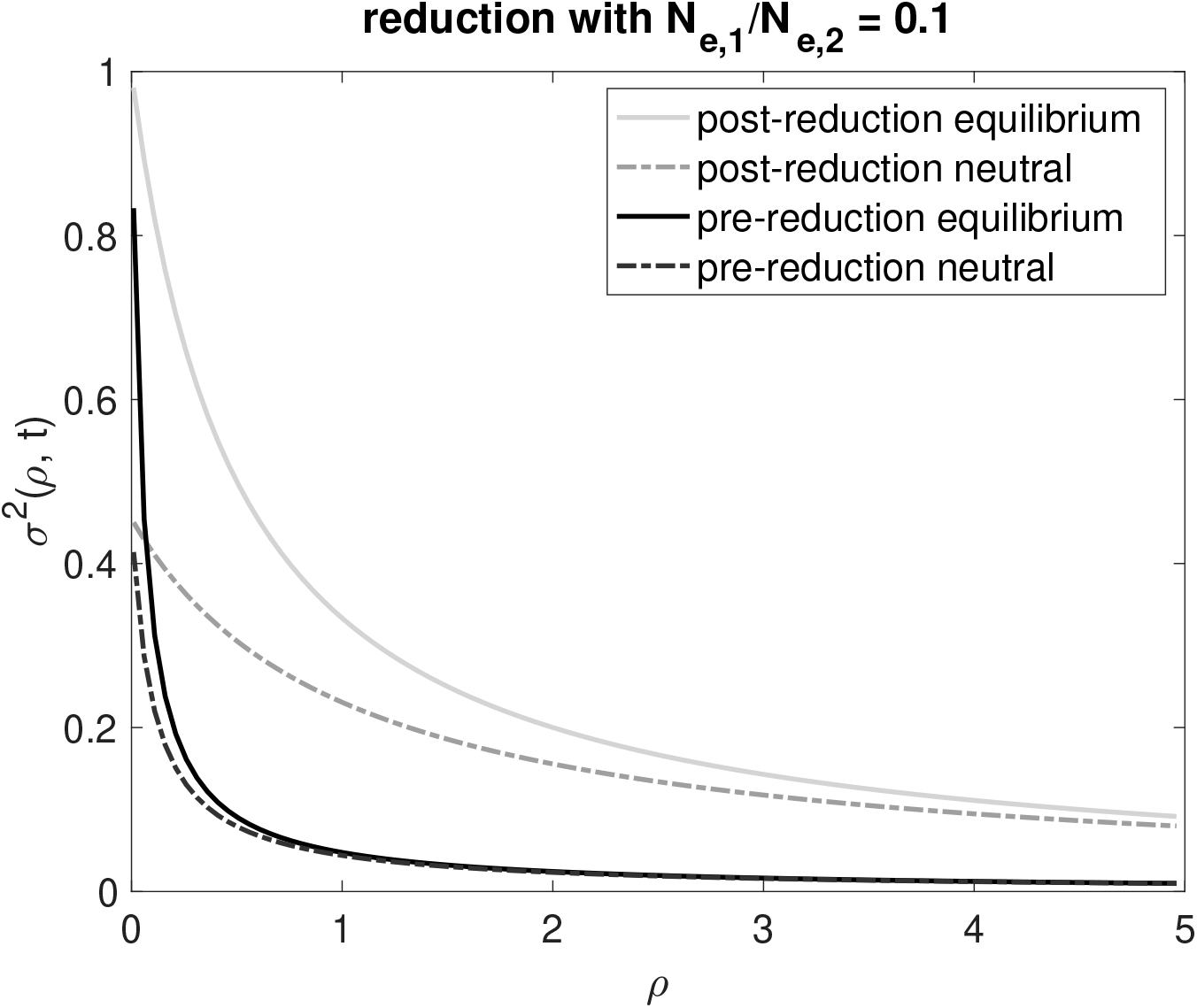
LD levels before and after population size reduction. The model is the same as that used in Figure 5d.

**Figure S6:**
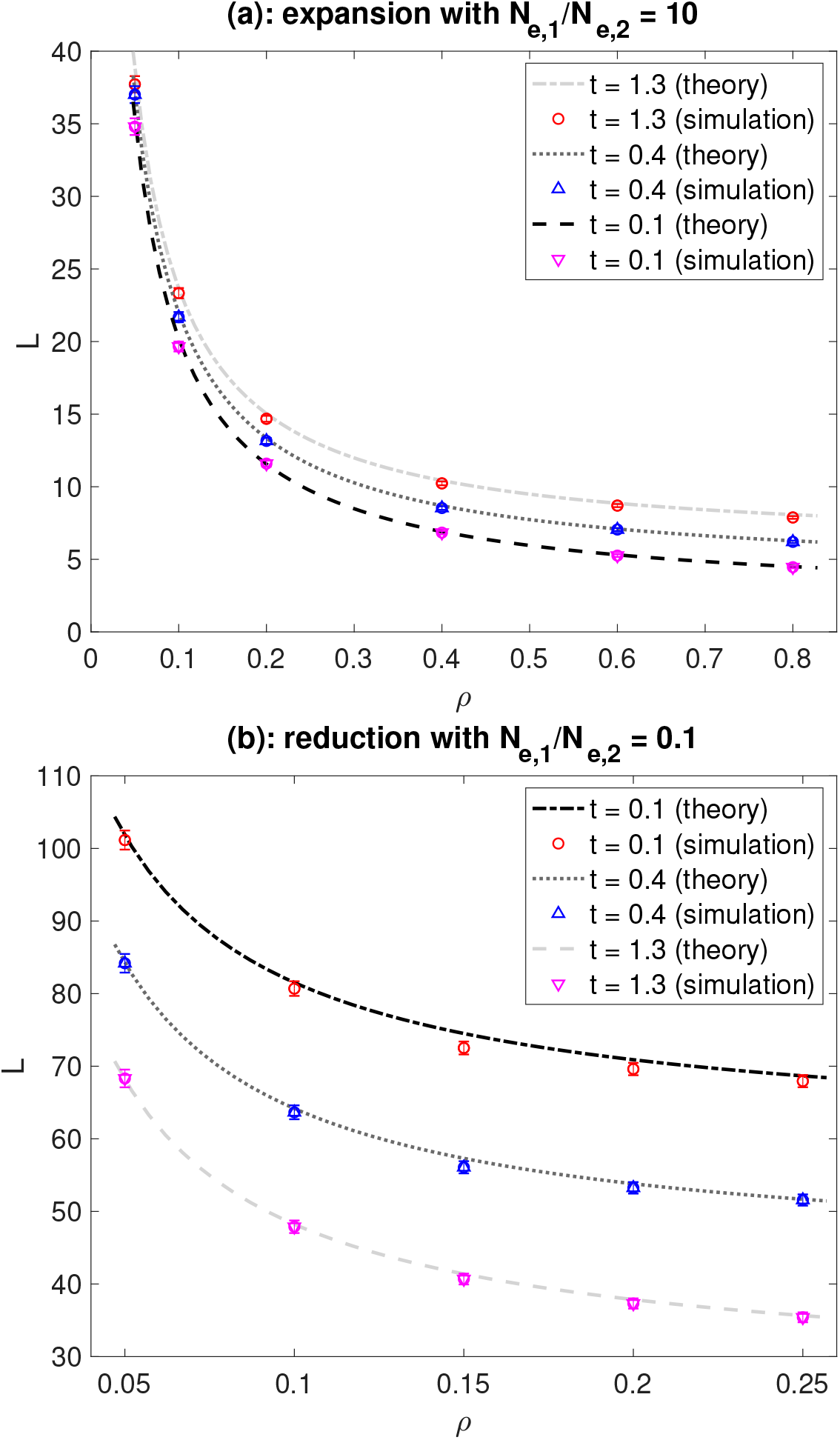
Expected total branch length *L* with demographic changes and strong balancing selection. We used the method described in the main text and Supplementary Text S.6 to carry out coalescent simulations with stochastic allele frequency trajectories, and compared the results to the theoretical predictions obtained under deterministic allele frequency trajectories. The population experienced a one-step change in population size at time *t* before the present. The population size in the present and ancestral epochs are *N*_*e*,1_ and *N*_e,2_, respectively. Time (*t*) and the recombination frequency (ρ) are expressed in units of 2*N*_*e*,1_ generations. For the expansion model in (a), *N*_*e*,1_ = 20, 000 and *N*_*e*,2_ = 2, 000. For the reduction model in (b), *N*_*e*,1_ = 2, 000 and *N*_*e*,2_ = 20, 000. These are the same as those considered in Figure 5. The fitnesses of *A*_1_*A*_1_, *A*_1_*A*_2_, and *A*_2_*A*_2_ are *w*_11_ = 1 –*s*, *w*_22_ = 1, and *w*_22_ = 1 –*s*, where *s* = 0.0625. We have 2 min (*N*_*e*,1_, *N*_*e*,2_)*s* = 250, implying that selection is strong. The equilibrium frequencies of *A*_1_ and *A*_2_ are 0.5. The sample size is *n* = 20. The mutation rate from *A*_1_ to *A*_2_ is 6.25 × 10^-8^per generation, and the rate in the opposite direction is the same. For each parameter combination, 10,000 simulation replicates were conducted to obtain an estimate of *L*. The whiskers show *L* ± 2 ×(standard error).

**Figure S7:**
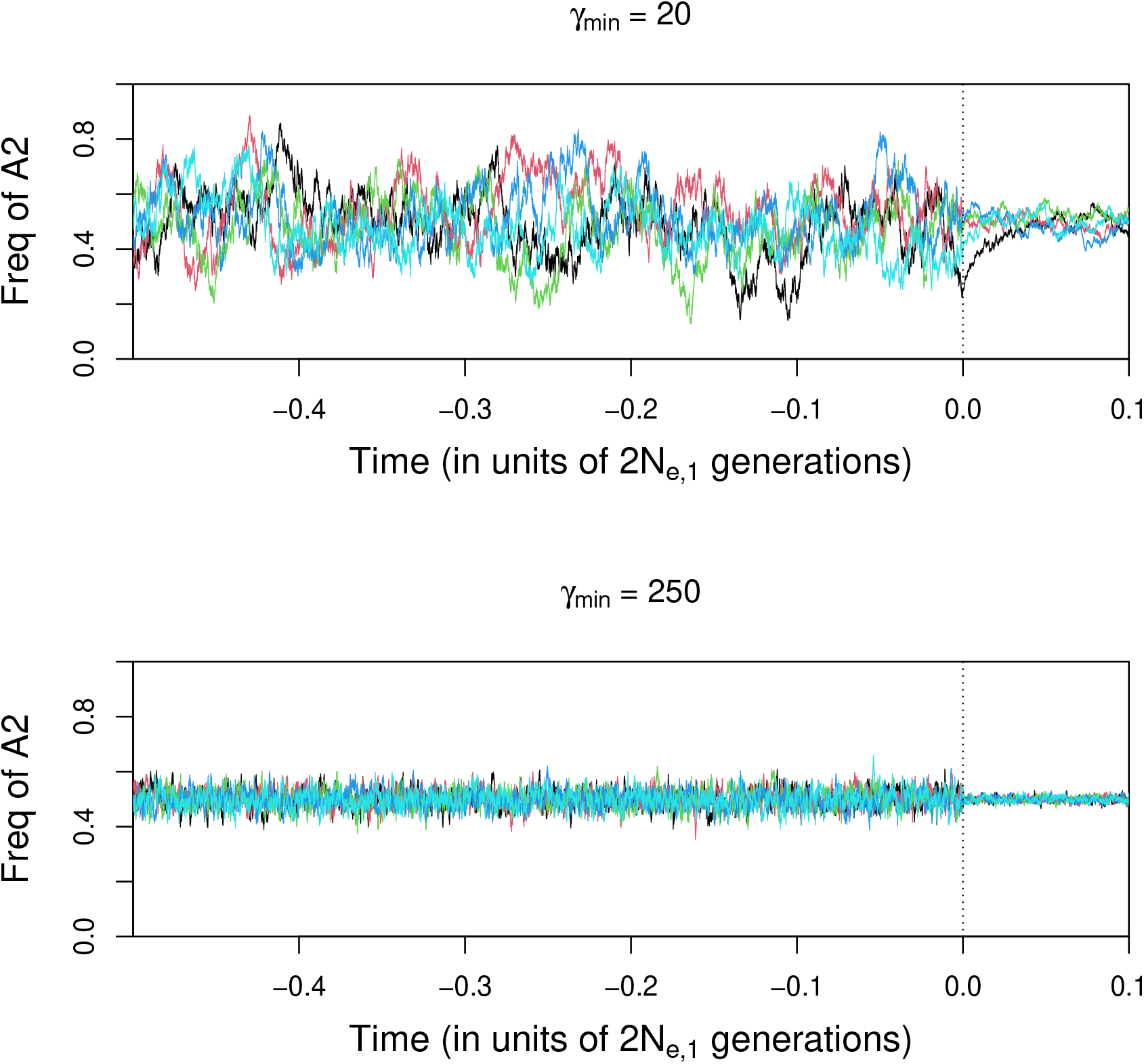
The effects of selection intensity on the extent of stochastic fluctuation in allele frequencies at the selected locus. We assumed the same population expansion model used in Figure S6, i.e., *N*_*e*,1_ = 20,000 (the current effective population size) and *N*_*e*,2_ = 2,000 (the effective population size in the ancestral epoch). The vertical dotted line indicates *t* = 0, which is when the population size increase takes place. The fitnesses of *A*_1_*A*_1_, *A*_1_*A*_2_, and *A*_2_*A*_2_ are *w*_11_ = 1 –*s, w*_12_ = 1, and *w*_22_ = 1 –*s*. Selection intensity is measured by *γ_min_* = 2*N_e,min_s*, where *N_e,min_* = min(*N*_*e*,1_, *N*_*e*,2_). The mutation rate from *A*_1_ to *A*_2_ is 6.25 ×10^-8^ per generation, and the rate in the opposite direction is the same. For each parameter combination, allele frequency trajectories of *A*_2_ were obtained using the forward simulation methods detailed in Supplementary Text S.6. In each plot, the trajectories from five simulations are included, and are shown in different colours.

**Figure S8:**
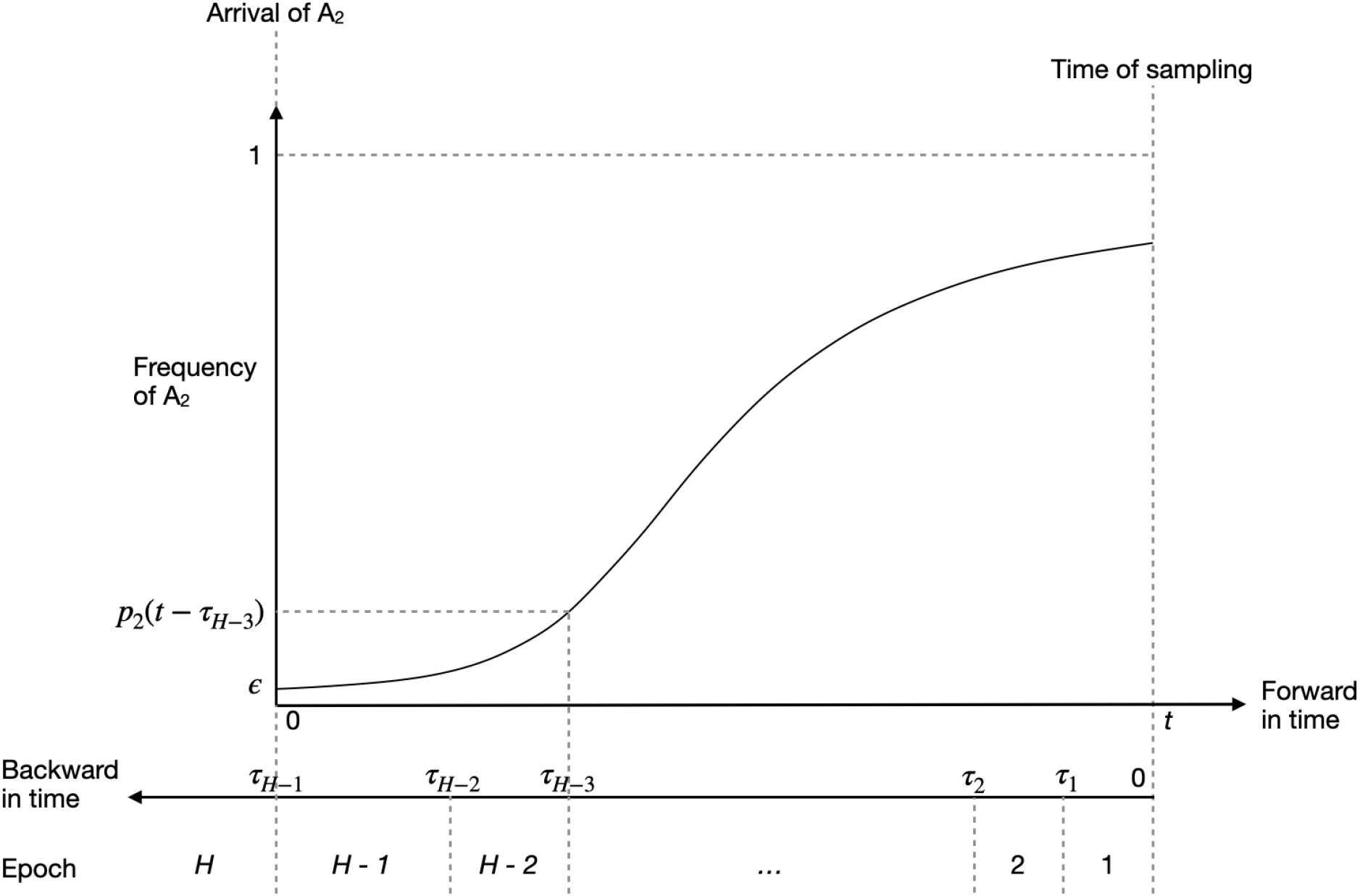
A diagram showing the discretisation scheme used to obtain the expected total branch length and the site frequency spectrum under the model of recent balanced polymorphism.

**Figure S9:**
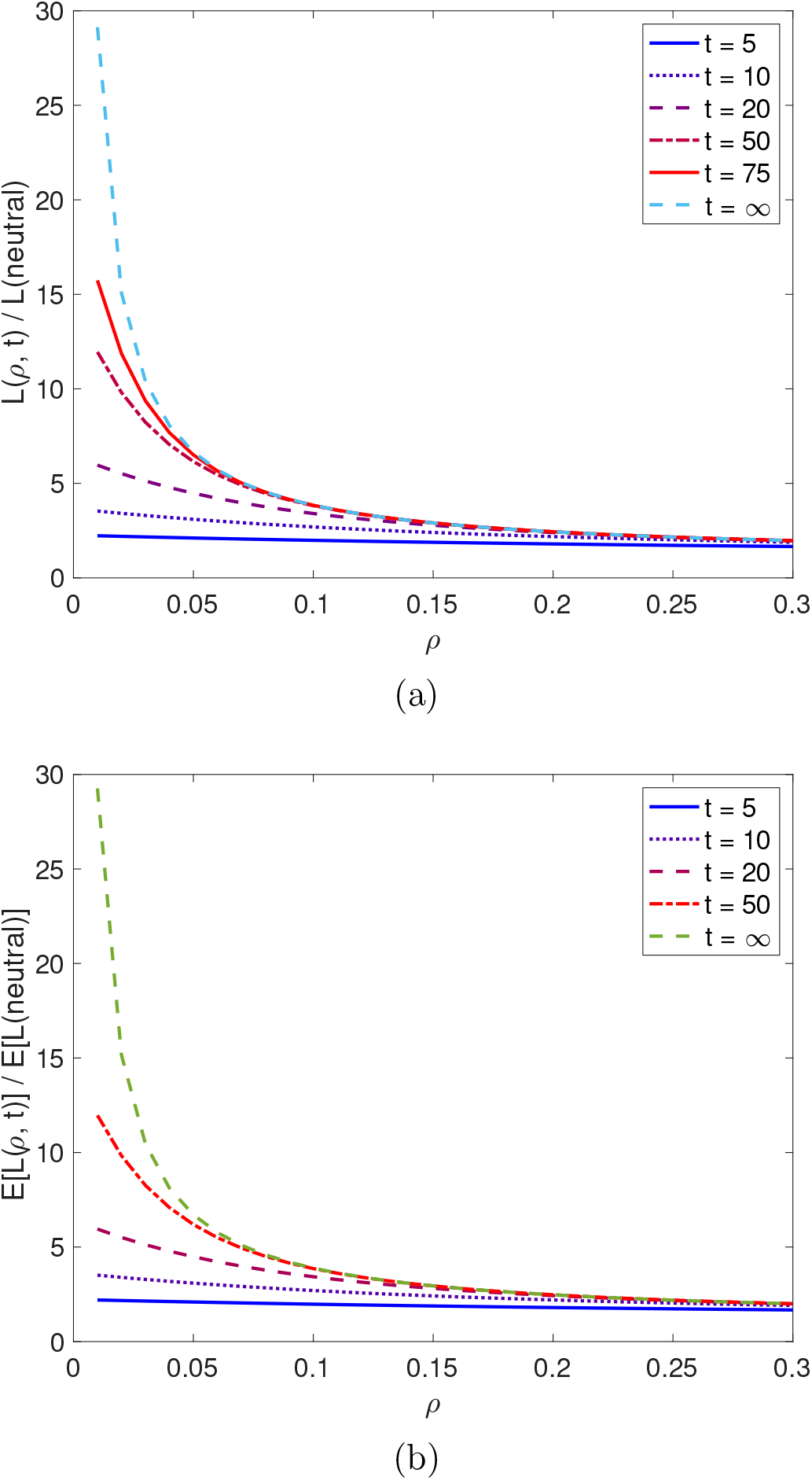
The approach to the equilibrium diversity level. The parameters are the same as those used in Figures 6 and 7. The sample size is 20. 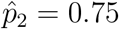 in (a) and 0.5 in (b). Note that the curves are based on a model without reversible mutation between the two selected variants *A*_1_ and *A*_2_. They overestimate the increase in diversity when *ρ* is very small.

**Figure S10:**
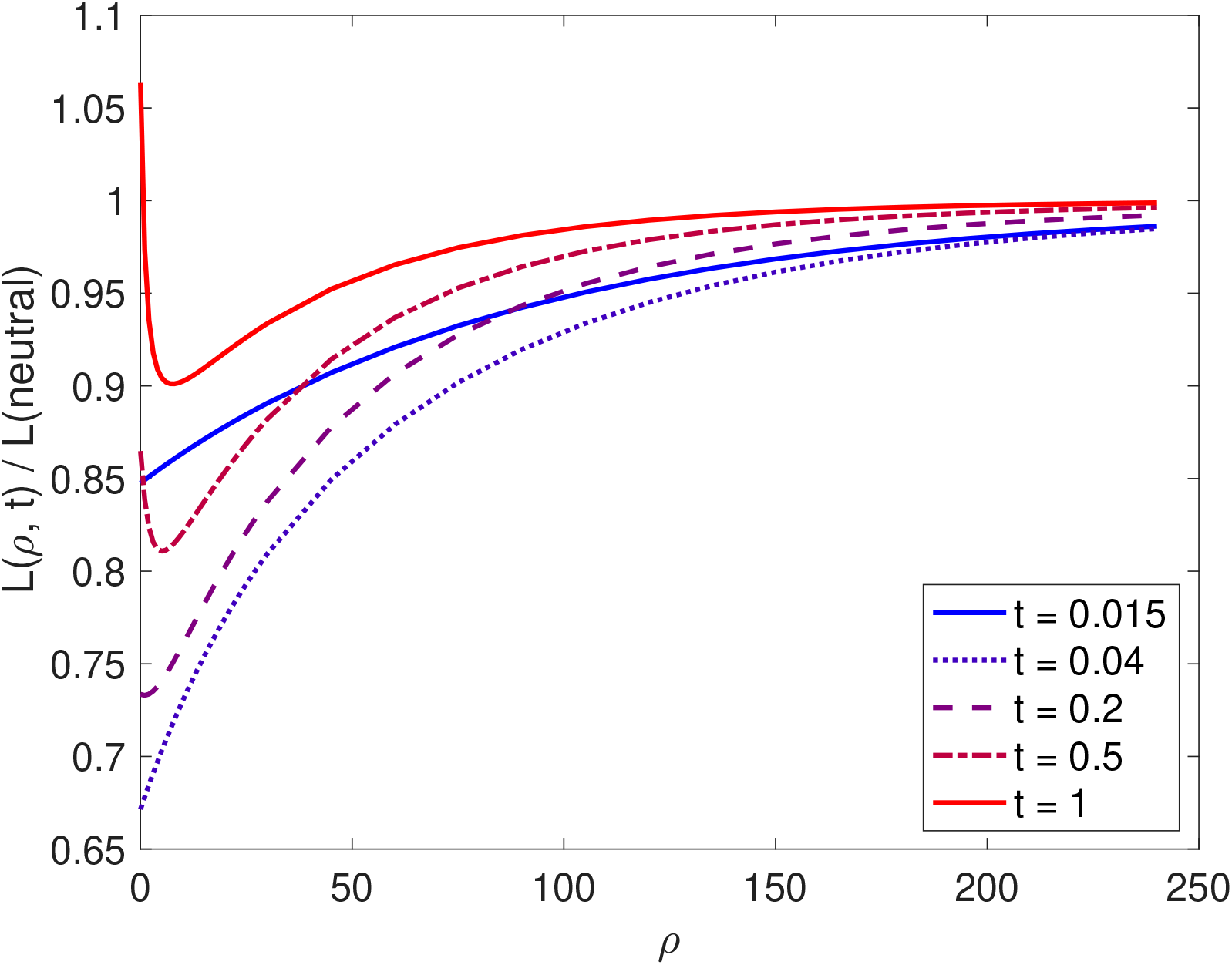
Neutral diversity in genomic regions surrounding a recently-emerged variant under balancing selection. The parameters are the same as in Figure 7 in the main text, except that the sample size is *n* = 20.

**Figure S11:**
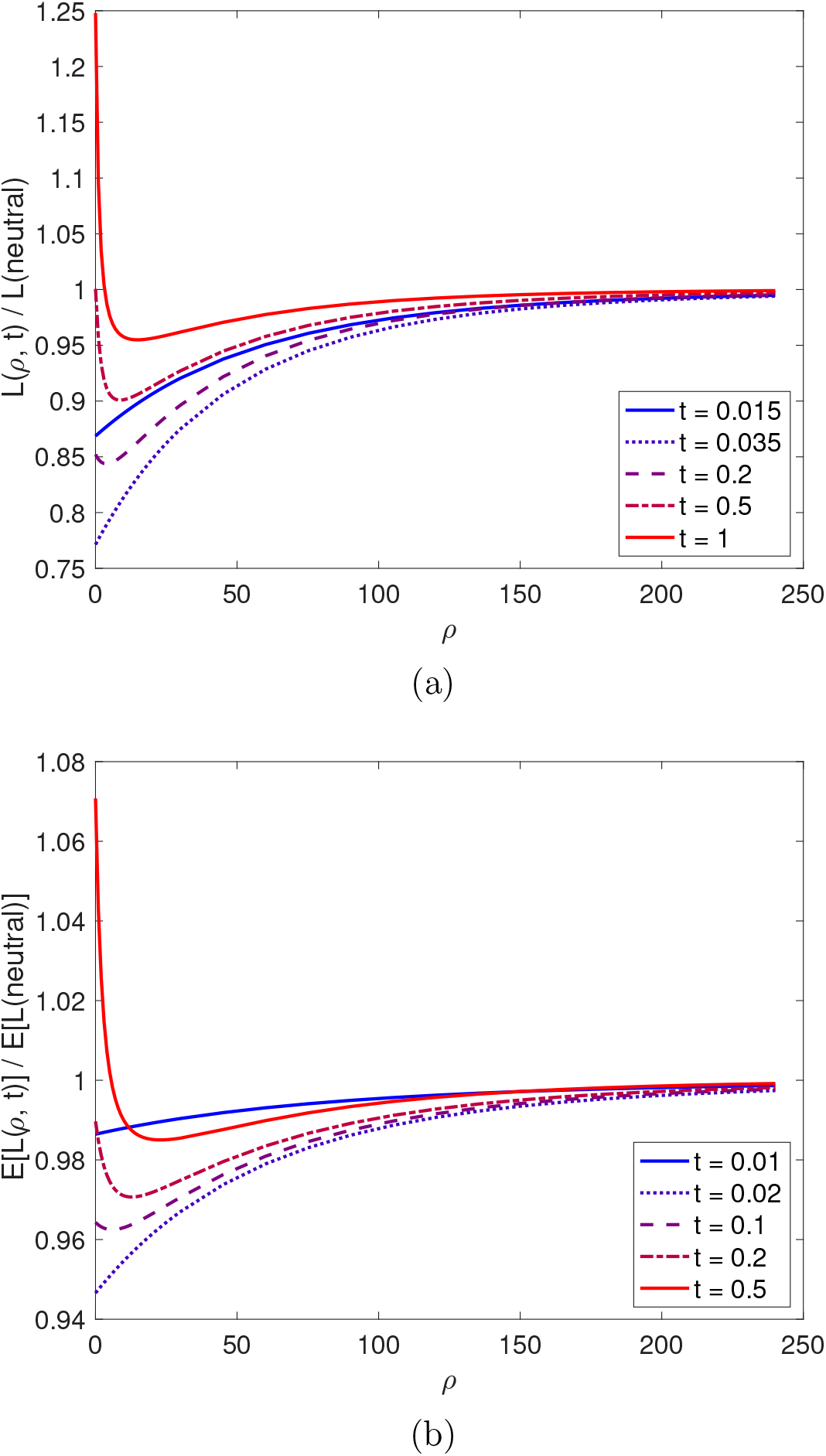
Neutral diversity level in genomic regions surrounding a recently-emerged balanced polymorphism. These figures are analogous to that in Figure 7, except that in (a) 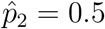 and in (b) 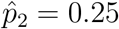. The sample size is two.

**Figure S12:**
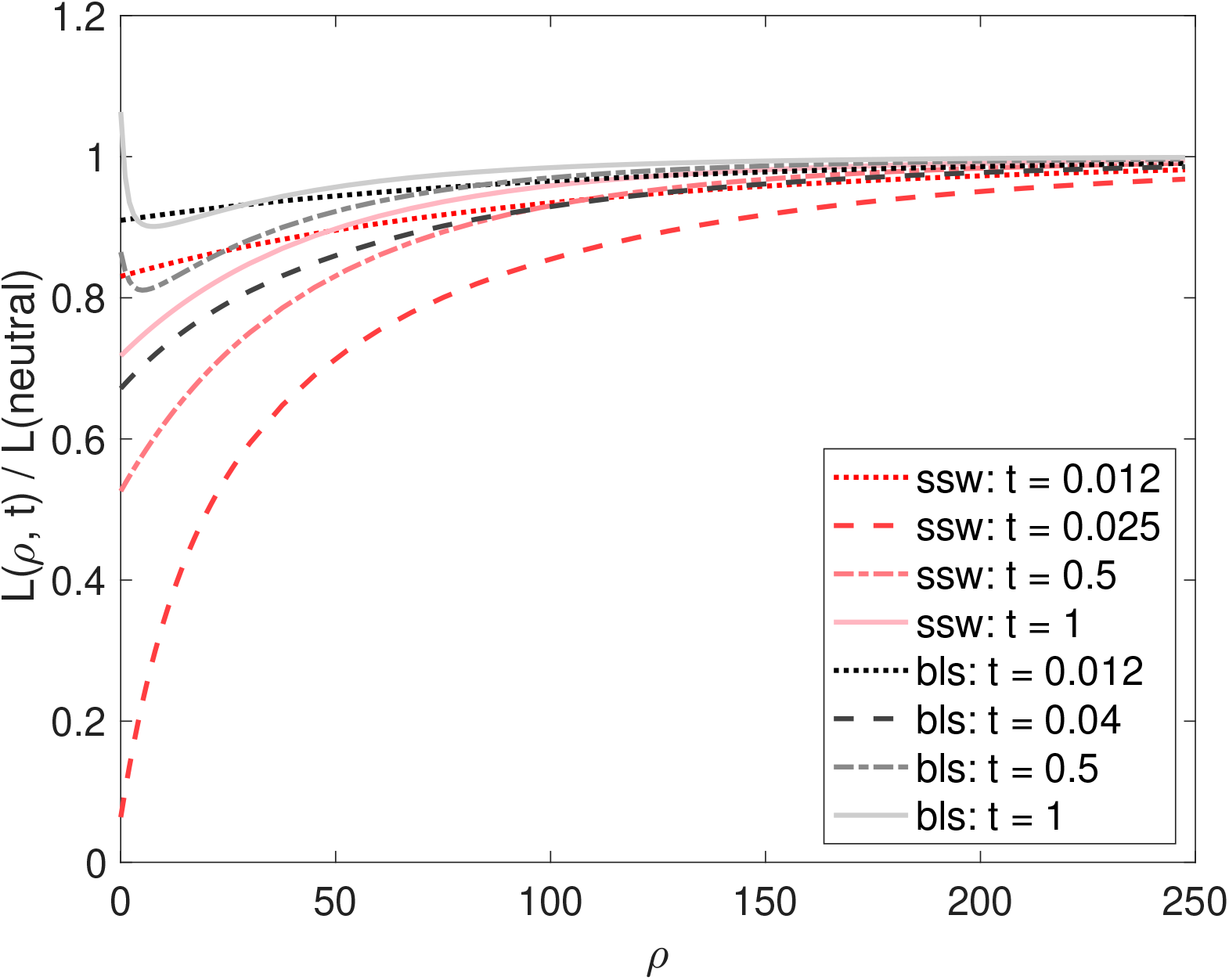
Comparing recent balancing selection with the corresponding sweep model with respect to their effects on diversity levels in surrounding genomic regions. The models and their parameters are the same as those in Figure 8, expect that *n* = 20.

**Figure S13:**
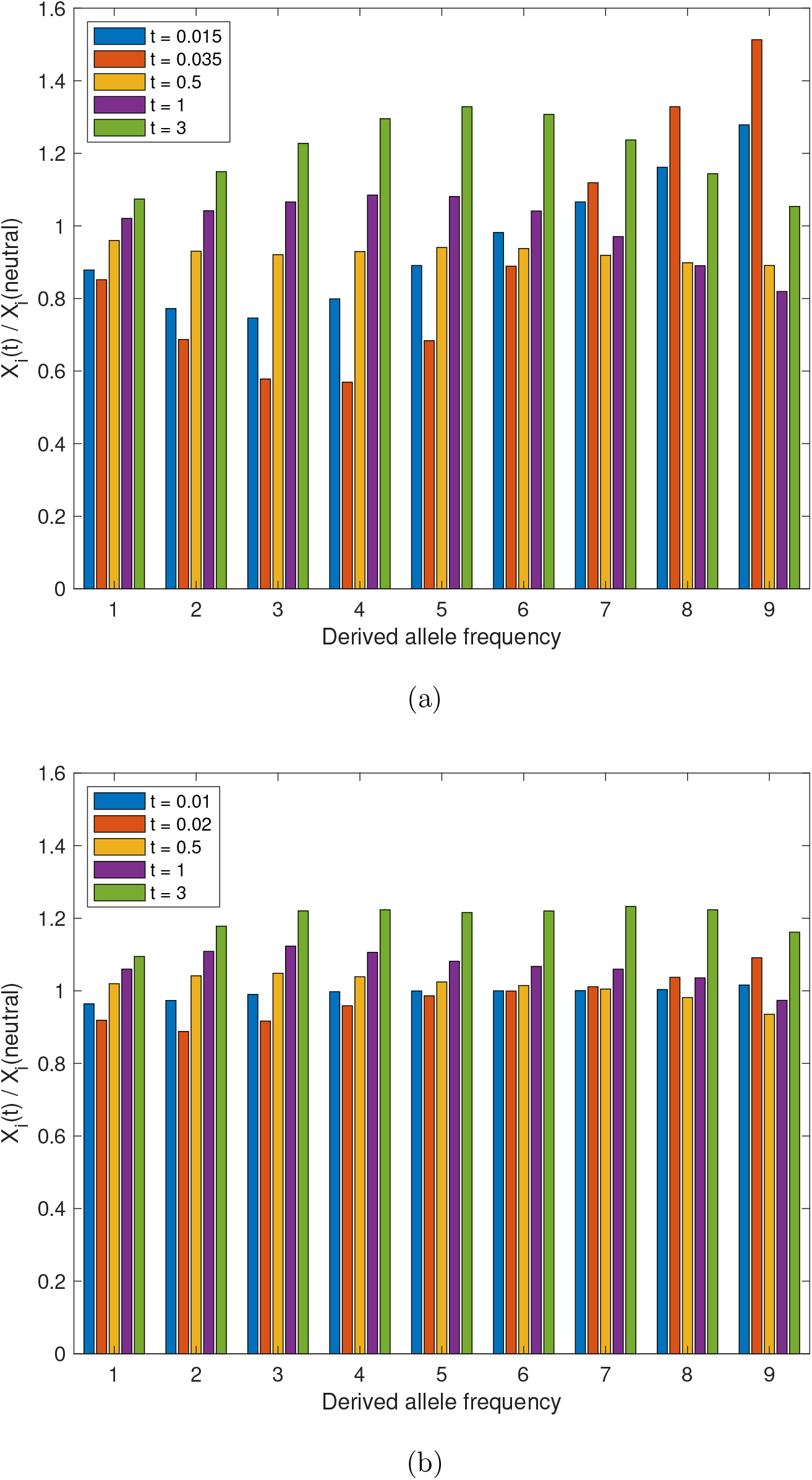
The SFS for the balancing selection models considered in Figure S11. In (a) 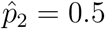 and in (b) 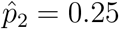.

**Figure S14:**
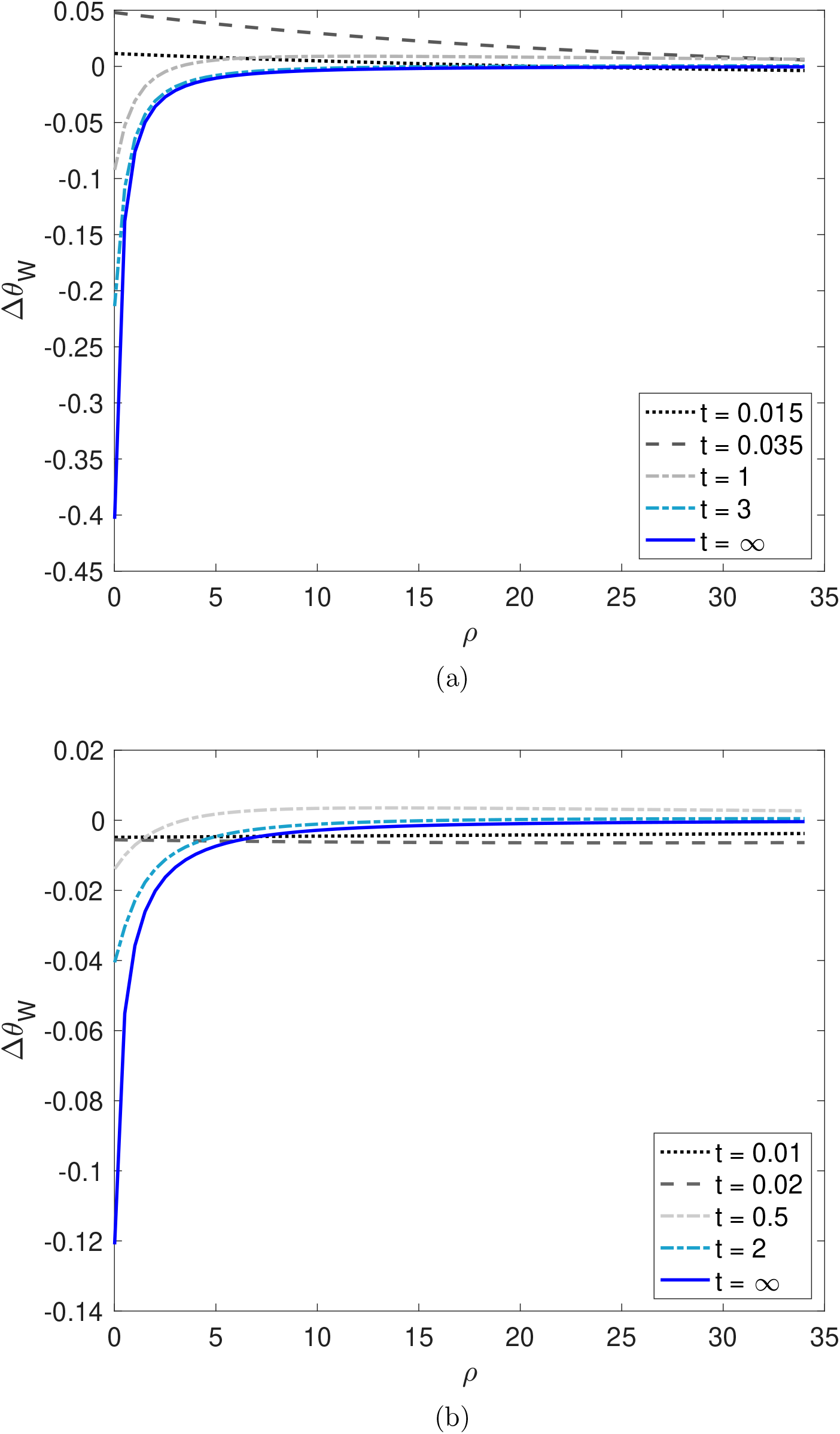
Δ*θ_W_* as a function of *ρ* and *t* for the balancing selection models considered in Figure S11. The sample size is 10. In (a) 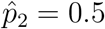 and in (b) 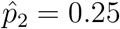.

**Figure S15:**
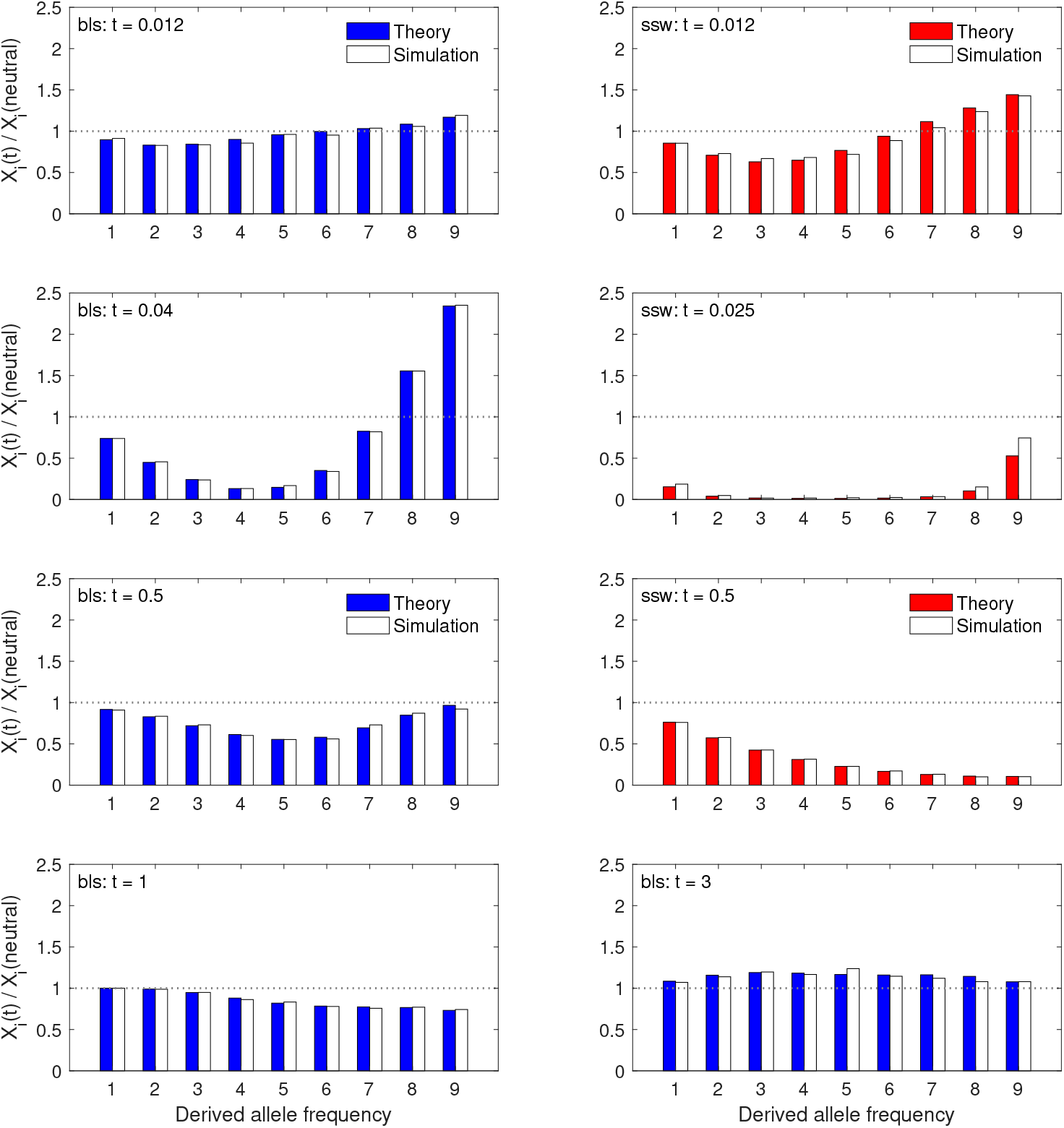
The SFS at various time points after the arrival of the selected variant for a sample of 10 alleles. The balancing selection (bls) and selective sweep (ssw) models are the same as those shown in Figure 9 in the main text (i.e., *γ*_1_ = 500 and 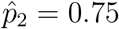). The scaled recombination frequency between the focal neutral site and the selected site is *ρ* = 2.

**Figure S16:**
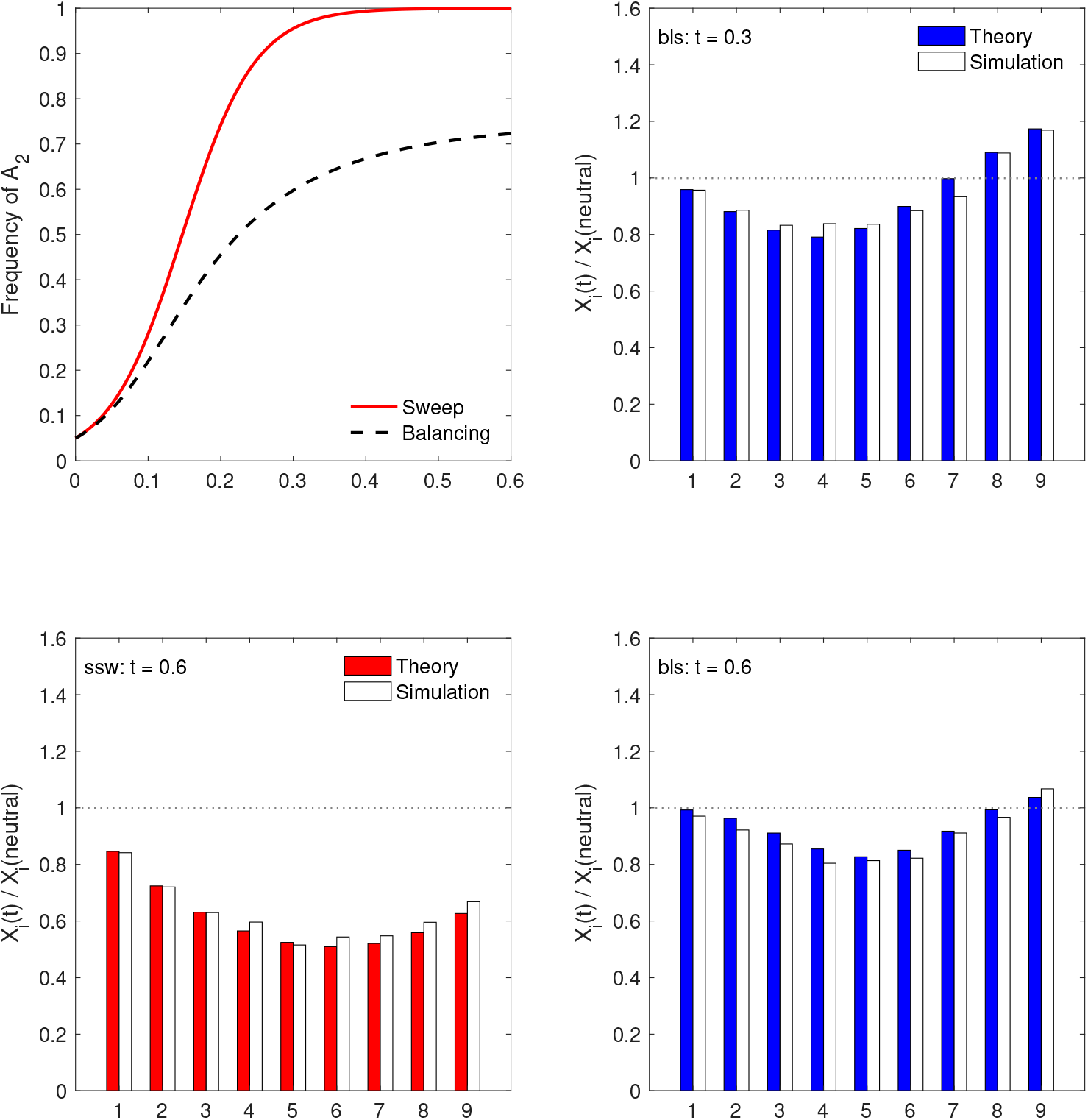
The SFS at various time points after the arrival of the selected variant for a sample of 10 alleles. The balancing selection (bls) model has *γ*_1_ = 20 and 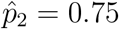, and the corresponding sweep (ssw) model was also simulated. The scaled recombination frequency between the focal neutral site and the selected site is *ρ* = 2.

